# The gut microbiome promotes mitochondrial respiration in the brain of a Parkinson’s disease mouse model

**DOI:** 10.1101/2024.12.18.629251

**Authors:** Livia H. Morais, Linsey Stiles, Milla Freeman, Anastasiya D. Oguienko, Jonathan D. Hoang, Jeff Jones, Baiyi Quan, Jack Devine, Justin S. Bois, Tsui-Fen Chou, Joanne Trinh, Martin Picard, Viviana Gradinaru, Sarkis K. Mazmanian

**Affiliations:** Division of Biology & Biological Engineering, California Institute of Technology, Pasadena, CA, USA; Aligning Science Across Parkinson’s (ASAP) Collaborative Research Network, Chevy Chase, MD 20815; UCLA Metabolomics Center, David Geffen School of Medicine, University of California, Los Angeles, CA, USA; Institute of Neurogenetics, University of Lübeck, 23538, Lübeck, Germany; Departments of Psychiatry and Neurology, Columbia University Irving Medical Center, New York, NY, USA; Departments of Medicine and Endocrinology, David Geffen School of Medicine, University of California, Los Angeles, CA, USA

## Abstract

The pathophysiology of Parkinson’s disease (PD) involves gene-environment interactions that impair various cellular processes such as autophagy, lysosomal function, or mitochondrial dysfunction. Specifically, mitochondria-associated gene mutations increase PD risk, mitochondrial respiration is altered in the PD brain, and mitochondrial-damaging toxins cause PD-like motor and gastrointestinal symptoms in animal models. The gut microbiome is altered in PD patients and represents an environmental risk, however a relationship between mitochondrial function and the microbiome in PD has not been previously established. Herein, we report that striatal mitochondria are functionally overactive in α-synuclein-overexpressing (ASO) mice, a model of PD, and that microbiome depletion restores respiration and mitochondria-associated gene expression patterns to wild-type levels. ASO mice harboring a complex microbiome produce increased reactive oxygen species in the striatum whereas germ-free counterparts express elevated levels of antioxidant proteins that may buffer against oxidative damage. Indeed, antioxidant treatment improves motor performance in ASO mice and, remarkably, blocking oxidant scavenging in germ-free mice induces α-synuclein-dependent motor deficits. Thus, the gut microbiome increases mitochondrial respiration and oxidative stress in the brain, which enhances motor symptoms in a mouse model of PD.

## INTRODUCTION

Parkinson’s disease (PD) is the second most common neurodegenerative disease in the United States, affecting 1% of the population over the age of 60, with often debilitating motor symptoms such as rigidity, tremors, and postural instability. Aggregation of the neuronal protein α-synuclein (αSyn) is believed to cause PD via death of dopaminergic neurons of the nigrostriatal pathway in the brain (*1*). While disease etiology is incompletely understood, there is substantial evidence for involvement of impaired mitochondrial function in PD. Mutations in several genes encoding mitochondria-associated proteins (*PINK1*, *PARK2*, and *PARK7*) are strongly linked to familial forms of PD (*2*). Alterations in mitochondrial respiration, dynamics, and quality control mechanisms are common in PD and its associated animal models (*3–5*), with accompanying increases in oxidative stress (*6*). Mitochondrial respiratory capacity is reduced in tissue extracts from the substantia nigra (*7*, *8*), a brain region associated with PD (*9*), and patients display elevated mitochondrial DNA mutations (*5*, *10*). Toxins that inhibit mitochondrial respiration cause neurodegeneration and motor symptoms in rodents and non-human primates (*11–14*), and mice harboring human mutations in mitochondria-associated genes recapitulate PD-like outcomes (*15–17*). Accumulation and aggregation of αSyn leads to impaired mitochondrial protein import systems, as well as altered mitochondrial dynamics and bioenergetics (*18–22*). Further, neuronal vulnerability and neurodegeneration in PD is associated with high mitochondrial energy demand and increased oxidative stress in certain cell types including dopaminergic neurons (*23*).

Although predominantly viewed as a brain disorder, many PD patients experience considerable non-motor symptoms including sleep disturbances, hyposmia, and gastrointestinal (GI) issues such as constipation, gastroparesis, and abdominal pain that usually manifest many years before a PD diagnosis (*24*, *25*). Braak’s hypothesis proposes that αSyn pathology may initiate in the GI tract and eventually reach the brain stem, substantia nigra, and neocortex via the vagus nerve (*26*). In support of this model, injection of αSyn fibrils into the intestines of mice and rats results in gut symptoms and, over time, brain neurodegeneration and motor deficits (*27*). In some studies, severing the vagus nerve in animals halts progression of pathology into the brain (*28*), and epidemiologic data suggest that vagotomies are protective against the development of PD in humans (*29*). Several studies in mice revealed that intestinal inflammation exacerbates PD-like symptoms (*30–32*), which corroborates findings in humans that inflammatory bowel disease may be a risk factor for PD (*33*). Accordingly, some forms of PD may originate in the periphery (body-first) and others in the brain (brain-first) (*34*, *35*).

Differences in gut microbiome composition between PD patients and household or matched population controls have been reproduced in numerous cohorts (*36–41*), with decreased abundance of health-promoting microbial species and increased pro-inflammatory bacterial taxa in the PD microbiome. Further, the microbiome profoundly impacts motor performance, gut function, and αSyn pathology in multiple PD mouse models (*20*, *30*, *42–46*), and pathogenic gut bacterial species accelerate disease outcomes (*47*, *48*) while antibiotic treatment or germ-free status improves motor symptoms in animals (*48–50*). Interestingly, fecal microbiota transplants (FMT) from PD patient donors into αSyn-overexpressing (ASO) mice results in worse motor performance compared to microbiomes from non-disease donors (*48*). Clinical studies that restore microbiome profiles to PD patients via FMT from healthy donors have shown promise in early stage human trials (*51–53*), with additional microbiome-based treatments in clinical development (*54*). Thus, an altered microbiome may represent both a contributor to PD etiology and a potential target for new interventions.

Herein, we explored the hypothesis that the gut microbiome impacts motor symptoms in ASO mice via modulation of mitochondrial function in the brain. We report that enhanced mitochondrial respiration, elevated levels of reactive oxygen species (ROS), αSyn-dependent brain pathology, and motor deficits in ASO mice are dependent on a complex microbiome.

Interestingly, mitochondria in the striatum of germ-free ASO mice, which do not display motor symptoms, produce increased levels of antioxidant proteins that neutralize ROS and pharmacologic disruption of redox homeostasis promotes motor defects in ASO mice lacking a microbiota. Thus, gut microbial stimulation of mitochondrial respiration and oxidative stress in the brain enhances motor symptoms in αSyn overexpressing mice, a process which may be implicated in subsets of PD patients with mitochondrial dysfunction.

## RESULTS

### The microbiome regulates motor behavior and gene expression in the brains of ASO mice

ASO mice are an experimental model for synucleinopathies, including PD, exhibiting robust motor symptoms and αSyn pathology in brain regions linked to human disease (*55*). We previously implicated a critical role for the microbiome in this animal model, reporting that ASO mice reared under germ-free (GF) conditions do not display motor deficits, neuroinflammation, or brain pathology, unlike mice with a standard laboratory microbiome (SPF; specific pathogen-free) (*48*). Here, we confirm that at four months of age, ASO-GF mice perform similarly to WT animals in the challenging beam and pole descent tests, as measured by the time to cross a beam or the time to climb down a pole, respectively (**Fig. 1, A and B**). We observed no significant differences between WT-SPF and WT-GF mice. Statistical analysis with a mixture model confirmed that a significantly larger fraction of ASO-GF mice successfully crossed the beam and descended the pole compared to ASO-SPF mice (**Fig. 1, A and B, lower panels**). Therefore the microbiome is required for motor deficits in mice that overexpress αSyn, similar to other preclinical models of PD (*49, 56–59*).

**Figure 1.**
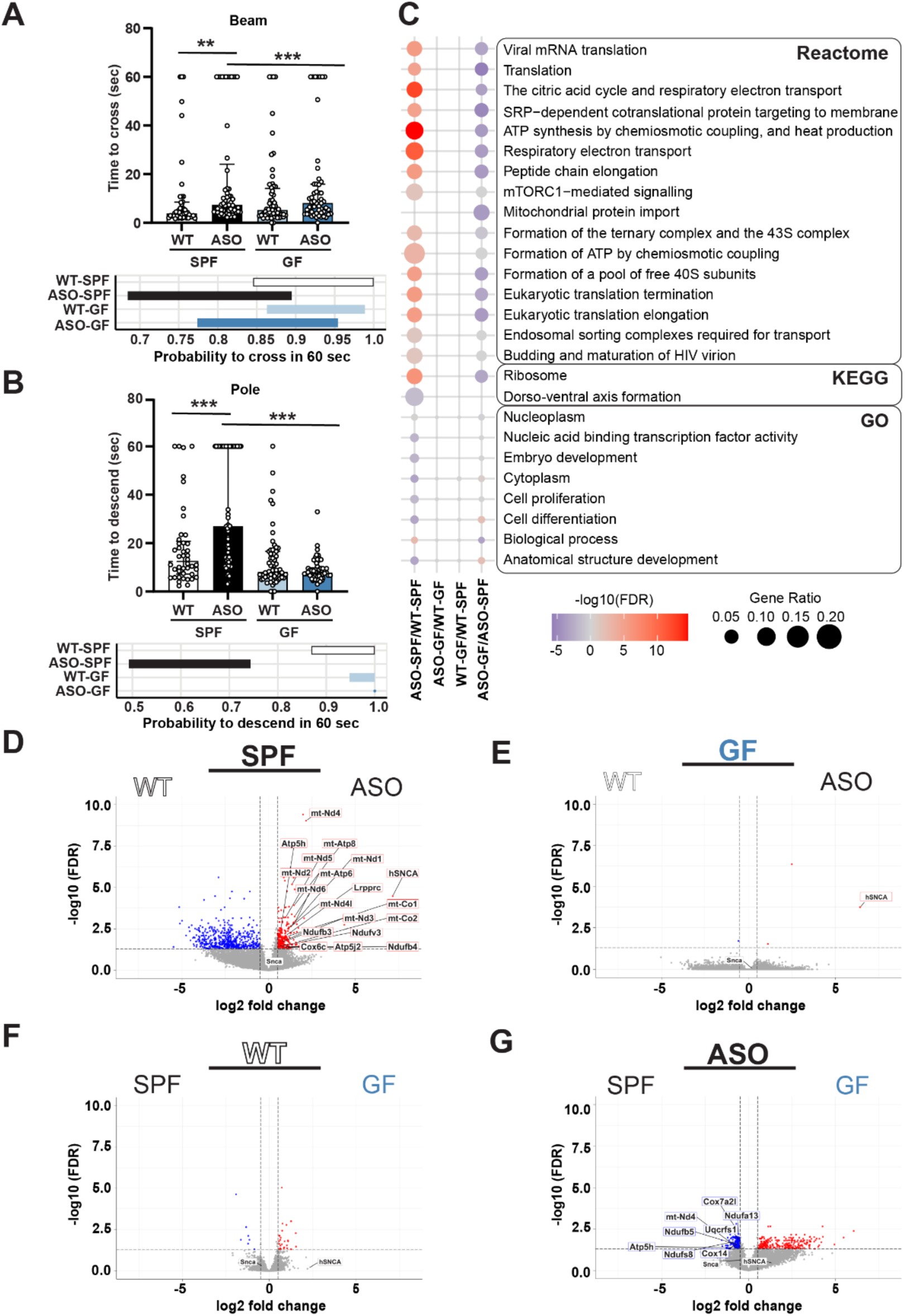
The gut microbiome regulates motor function and genes involved in mitochondrial respiration in ASO mice. (**A-B**) Motor testing (**A**) Top: time to cross challenging beam. Bottom: 95% confidence intervals (CIs) of probability to cross successfully within 60 seconds. (**B**) Top: time to descend pole. Bottom: 95% CIs of probability to descend successfully within 60 seconds. (**C-F**) Bulk quantitative RNA-sequencing of whole striatal tissue to assess differential gene expression between groups, comparing the effects of genotype (WT x ASO) and the gut microbiome (SPF x GF). (**C**) Gene set enrichment analysis (GSEA) conducted on upregulated and downregulated genes against KEGG, GO, and Reactome databases. The top 10 most significant terms for each comparison are included. (**D-G**) Volcano plots showing genotype effects (**D-E**) and microbiome effects (**F-G**). Only DEGs associated with citric acid cycle, respiratory electron transport, and ATP synthesis by chemiosmotic coupling and heat production are identified by protein name in D-G, as well as *Snca* (endogenous αSyn), and *hSNCA* (transgenic human αSyn). Dashed vertical lines show the thresholds of log_2_(fold change) ≥ 0.5 and ≤ −0.5. SPF, specific pathogen-free; GF, germ-free; WT, wild-type; ASO, Thy1-α-synuclein overexpressing. **Statistical details:** (**A-B**) Bar plots show time to descend the pole/cross the beam. Data are shown as median with interquartile range. Pairwise comparisons were calculated with Kruskal-Wallis test followed by Conover’s post-hoc test. Beam traversal and pole descent times were analyzed with a generative mixture model (y ∼ ωLogNormal(μ, σ) + (1 - ω)δ_y60_). CIs were calculated using maximum likelihood estimation (MLE) followed by parametric bootstrap. The ω 95% CI parameter describes the probability to descend the pole/cross the beam within 60 sec. Significance: ** *p* < 0.01, *** *p* < 0.001. Sample size: n=9-13 mice per group, 1-3 trials per mouse. Data combined from 4 different cohorts. (**C-G**) Differentially expressed genes and overrepresented pathways were determined using a two-sided adjusted *p*-value with the Benjamini-Hochberg correction method. Significance: *q* < 0.05. Sample size: n=5 per group.

To explore functional consequences of gut microbiome depletion during αSyn overexpression, we performed RNA sequencing of whole striatal tissue, a brain region critically implicated in the ASO model and in human PD (*60–64*). ASO-SPF mice (the only group with motor deficits) displayed a distinct transcriptome compared to both WT groups that only partially overlapped with ASO-GF mice (**Fig. 1C**). Cell-type enrichment analysis of all differentially expressed genes (DEGs) revealed general overrepresentation of neuronal genes but no major differences in gene expression patterns between various cell subsets (**fig. S1A**). The total number of DEGs with significant adjusted *p*-values and log_2_ fold change (FC) > 0.5 varied across comparisons: 819 DEGs in ASO-SPF versus WT-SPF; 425 in ASO-SPF versus ASO-GF; 34 between WT-SPF and WT-GF; and 4 between ASO-GF and WT-GF (**Table S1**). Microbiome effects on transcription in the striatum of ASO animals, i.e., comparing ASO-SPF to ASO-GF mice, uncovered differences in gene expression programs linked to inflammation and metabolic activity (**fig. S1B, Table S1**). These results reveal that multiple gene expression programs in the striatum are activated by αSyn overexpression (ASO-SPF/WT-SPF) and are controlled by the microbiota (ASO-GF/ASO-SPF) (**Fig. 1C**).

### Gene expression related to mitochondrial respiration is controlled by gut microbiota

Gene set enrichment analyses (GSEA) on all upregulated and downregulated genes using the GO, KEGG, and Reactome databases (*65–67*) uncovered that ASO-SPF mice were altered in putative functions related to the citric acid cycle, respiratory electron transport, and ATP synthesis by chemiosmotic coupling and heat production compared to WT-SPF and ASO-GF counterparts (**Fig. 1C**). Thus, both gene and microbiome contributions shape energetic metabolism in the striatum of ASO mice. For genotype effects, DEGs that reached the significance criteria of q<0.05 and |log_2_(FC)| > 0.5 in the most highly upregulated pathways in ASO-SPF mice compared to WT-SPF mice (citric acid cycle, respiratory electron transport, and ATP synthesis by chemiosmotic coupling, and heat production) encode proteins of the mitochondrial electron transport chain (ETC), including Complex I (*mt-Nd1, mt-Nd2, mt-Nd3, mt-Nd4l, mt-Nd6, Ndufb3, Ndufb4, Ndufv3*) and Complex IV (*mt-Co1, mt-Co2, Cox6c*) (**Fig. 1D**, full gene names in **Table 1**). Importantly, these gene expression changes were absent when comparing WT and ASO mice under GF conditions (**Fig. 1E**). We also observed upregulation of genes associated with the respiratory electron transport pathway in ASO-SPF animals, including enrichment of integral components of the ATP synthase complex (*Atp5h, Atp5j2* and *mt-Atp8*) (**Fig. 1D**). For microbiome effects, ASO-SPF mice compared to ASO-GF were enriched in expression from genes for Complex I (*mt-Nd4, Ndufa13, Ndufb5, Ndufs8)*, Complex III (*Uqcrfs1, Cox7a2l, Cox14*) (**Fig. 1G**). Likewise, the ATP synthase complex, including ATP synthase membrane subunit d (*Atp5h*), was also enriched in the ASO-SPF group (**Fig. 1G**).

**Table 1.**
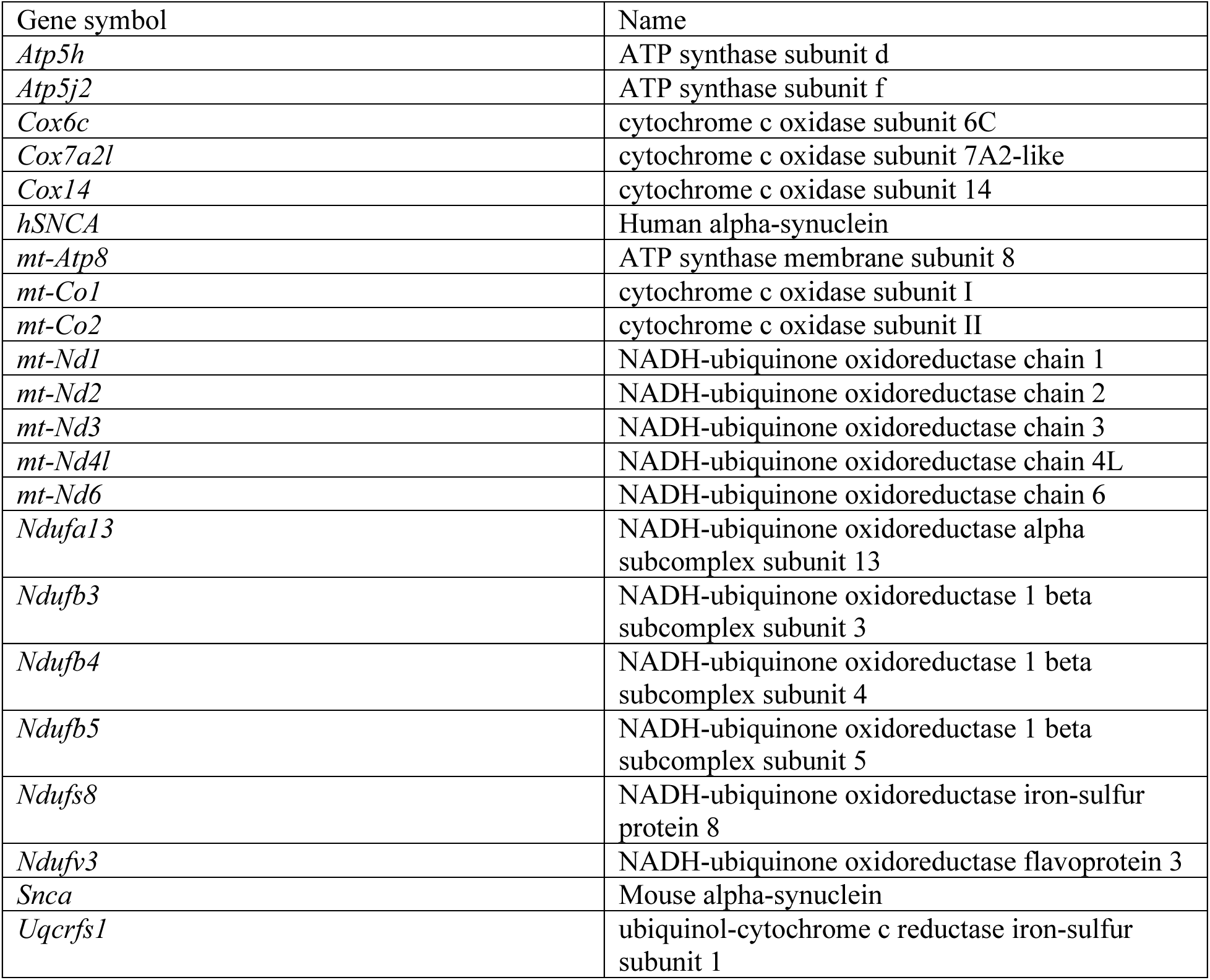
Genes symbols and names.

Comparison of WT-SPF and WT-GF animals did not reveal similar changes (**Fig. 1F**), demonstrating that alterations in mitochondrial-associated gene expression by the microbiome are dependent on αSyn overexpression. As expected, the αSyn-encoding transgene *hSNCA* was upregulated in ASO mice regardless of microbiome status, and *Snca* (the mouse αSyn gene) was not differentially regulated (**Fig. 1, E and F**). These findings determine that the microbiome robustly modulates mitochondrial respiration and energy metabolism in the striatum of ASO mice.

### Mitochondrial respiration in ASO mice is regulated by the microbiome

PD is associated with disruptions in oxidative phosphorylation (OxPhos), the primary cellular pathway for energy production (*68*). To validate whether mitochondrial energetic metabolism is influenced by the gut microbiome in ASO mice, we isolated mitochondria from the striatum and directly measured respiration, the series of biochemical reactions performed by Complexes I-V and other mitochondrial membrane proteins (*69*). When mitochondria were incubated with pyruvate and malate to drive Complex I respiration, ASO-SPF mice exhibited a robust increase in State 3u respiration (uncoupled respiration), with a trend towards increased State 3 respiration (ATP-linked respiration) (**Fig. 2, A and B**). Increased uncoupled respiration suggests enhanced capacity to transport and oxidize energy substrates in ASO-SPF mice compared to WT-SPF and both germ-free groups. We observed no significant differences in State 4o (proton leakage in the absence of ATP turnover) (**Fig. 2C**), State 2 (processes that consume membrane potential such as proton leakage and calcium cycling) (**fig. S2A**), or the respiratory control ratio (RCR) which estimates the capacity for substrate oxidation coupled to ATP synthesis (**fig. S2B**). Next, mitochondria were incubated with succinate to drive Complex II respiration, revealing increases in State 3u (**Fig. 2D**), State 3 (**Fig. 2E**), and State 4o (**Fig. 2F**) in ASO-SPF mice compared to the other animal groups. State 2 respiration and RCR were similar across groups in response to succinate (**fig. S2, C and D**). Conversely, mitochondria from ASO-GF mice exhibited similar oxygen consumption for Complex I and Complex II respiration as those from WT animals (**Figs. 2, A to F, S2, A to D**), confirming gene-microbiome interactions modulate mitochondrial respiration.

**Figure 2.**
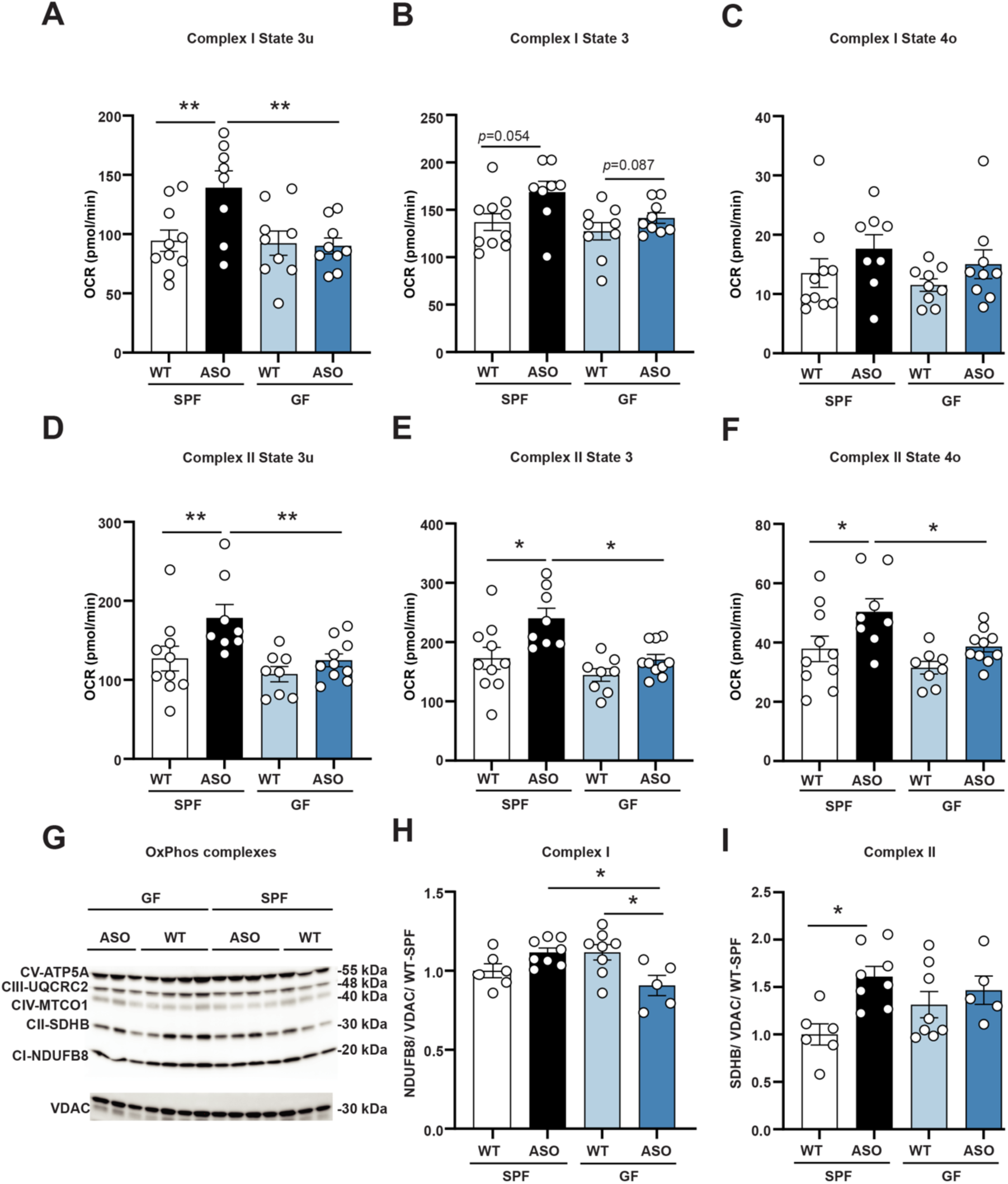
Gut microbiome regulates mitochondrial respiration in ASO mice. (**A-F**) Oxygen consumption rate (OCR) was measured in freshly isolated striatal mitochondria using Seahorse. (**A, D**) State 3u (uncoupled, maximal respiration). (**B, E**) State 3 (ATP-linked respiration). (**C, F**) State 4o (proton leak). (**G-I**) Western blotting for oxidative phosphorylation (OxPhos) complexes in isolated mitochondria. (**G**) Image of representative blot of two Western blots used for quantification showing levels of CV-ATP5A, CII-UQCRC2, CIV-MTCO1, CII-SDHB, and CI-NDUFB8. (**H**) Relative levels of Complex I NDUFB8 subunit in different groups. **I)** Complex II SDHB subunit. Protein levels were normalized to voltage-dependent anion channel 1 (VDAC1). SPF, specific pathogen-free; GF, germ-free; WT, wild-type; ASO, Thy1-α-synuclein overexpressing; CI, Complex I; CII, Complex II; CIII, Complex III; CIV, Complex IV; CV, Complex V; NDUFB8, oxidoreductase subunit B8; SDHB, succinate dehydrogenase subunit B; UQCRC2, ubiquinol-cytochrome c reductase core protein 2; MTCO1, cytochrome c oxidase subunit 1; ATP5A, ATP synthase subunit alpha. **Statistical details:** (**A-I**) Data were analyzed using a linear model (variable ∼ Genotype + Microbiome + Genotype*Microbiome) and pairwise comparisons with Benjamini-Hochberg (FDR) correction. Data are expressed as mean ± SEM. Significance: * *p* < 0.05, ** *p* < 0.01. (**A-F**) Sample size: n=8-10 per group, data combined from 4 different cohorts. (**G-I**) Sample size: n=6-8 per group, data combined from 2 different cohorts.

ETC activity is influenced by the abundance and assembly of OxPhos proteins (*70*). We measured levels of five OxPhos subunits in isolated mitochondria using Western blot analysis, which revealed that the gut microbiome influences expression of the Complex I protein NADH oxidoreductase subunit B8 (NDUFB8), with no significant difference between WT-GF and ASO-GF mice (**Fig. 2, G to I**). We observed no meaningful changes in levels of proteins related to Complex III [ubiquinol-cytochrome c reductase core protein 2 (UQCRC2)], Complex IV [cytochrome c oxidase subunit 1 (MTCO1)], or Complex V [ATP synthase subunit alpha (ATP5A)] (**fig. S2, E to G**). We further measured the activity of key enzymes in the respiratory chain and although not significant, NADH ubiquinone oxidoreductase (**fig. S3A**), succinate-ubiquinone oxidoreductase (**fig. S3B**), and cytochrome c oxidase (**fig. S3C**) trended toward increases in ASO-SPF animals compared to WT-SPF. We noted modest effects in Mitochondrial Health Index (MHI), which reflects mitochondrial specialization and respiratory capacity on a per-mitochondrion basis (*71, 72*), which was elevated in ASO-SPF animals compared to WT-SPF (**fig. S3D**), and the mitochondrial respiratory capacity (a normalized index similar to MHI) trended toward a slight increase in ASO-SPF vs. ASO-GF mice (**fig. S3E**).

To establish whether mitochondrial respiration and OxPhos protein expression are associated with increased mitochondrial biogenesis, we quantified the activity of citrate synthase (a Krebs cycle enzyme representing mitochondrial volume density), mitochondrial DNA (mtDNA) copy number (per cell) and mtDNA density (per mg of tissue) in total striatal extracts, and detected no differences between groups in any of these parameters (**fig. S3, F to H**). Next, we quantified expression of mitochondrial outer membrane markers [75-kDa glucose-regulated protein (GRP75) and the translocase of the outer mitochondrial membrane 20 (TOMM20)] and a mitochondrial nucleoid marker [mitochondrial transcription factor A (mtTFA)], none of which were affected by genotype or the gut microbiome (**fig. S3, J to L**). Together, these results show that increased mitochondrial respiration in ASO-SPF mice is likely not due to changes in mitochondrial abundance.

Overall, the gut microbiome increases mitochondrial activity in the striatum of 4-month-old ASO mice, which is prior to the onset of neurodegeneration in this model (*55*). We speculate that increased mitochondrial activity reflects either compensation for decreased bioenergetic efficiency or hypermetabolism due to αSyn overexpression, resulting in mitochondria working harder to meet energy demands within cells, as previously described in other contexts (*73–75*). While enhanced mitochondrial respiration by the microbiome may be part of the neurodegenerative process, more work is needed to determine if this activity contributes to neuronal loss.

### Microbial colonization increases oxidative stress in ASO mice

Altered mitochondrial respiration can manifest in various cellular and molecular outcomes, some of which can be captured by changes in mitochondria-associated proteins. Proteomic analysis of mitochondrial extracts isolated from the striatum revealed that mitochondrial protein expression is predominantly influenced by the gut microbiome, with remarkably minimal impact from genotype (**Fig. 3A, C and D**). Differentially expressed proteins most affected by the gut microbiome are implicated in pathways related to detoxification, ROS and glutathione metabolism, mitochondrial tRNA synthetases, protein import and sorting, and lipid metabolism (**Fig. 3B**). Notably, mitochondrial import and sorting proteins, essential for OxPhos complex biogenesis, including TIMM10 (translocase of inner mitochondrial membrane 10), TIMM9 (translocase of inner mitochondrial membrane 9), and TOMM22 (translocase of outer mitochondrial membrane 22) were enriched in SPF mice of both genotypes (**Fig. 3E** and **F**).

**Figure 3.**
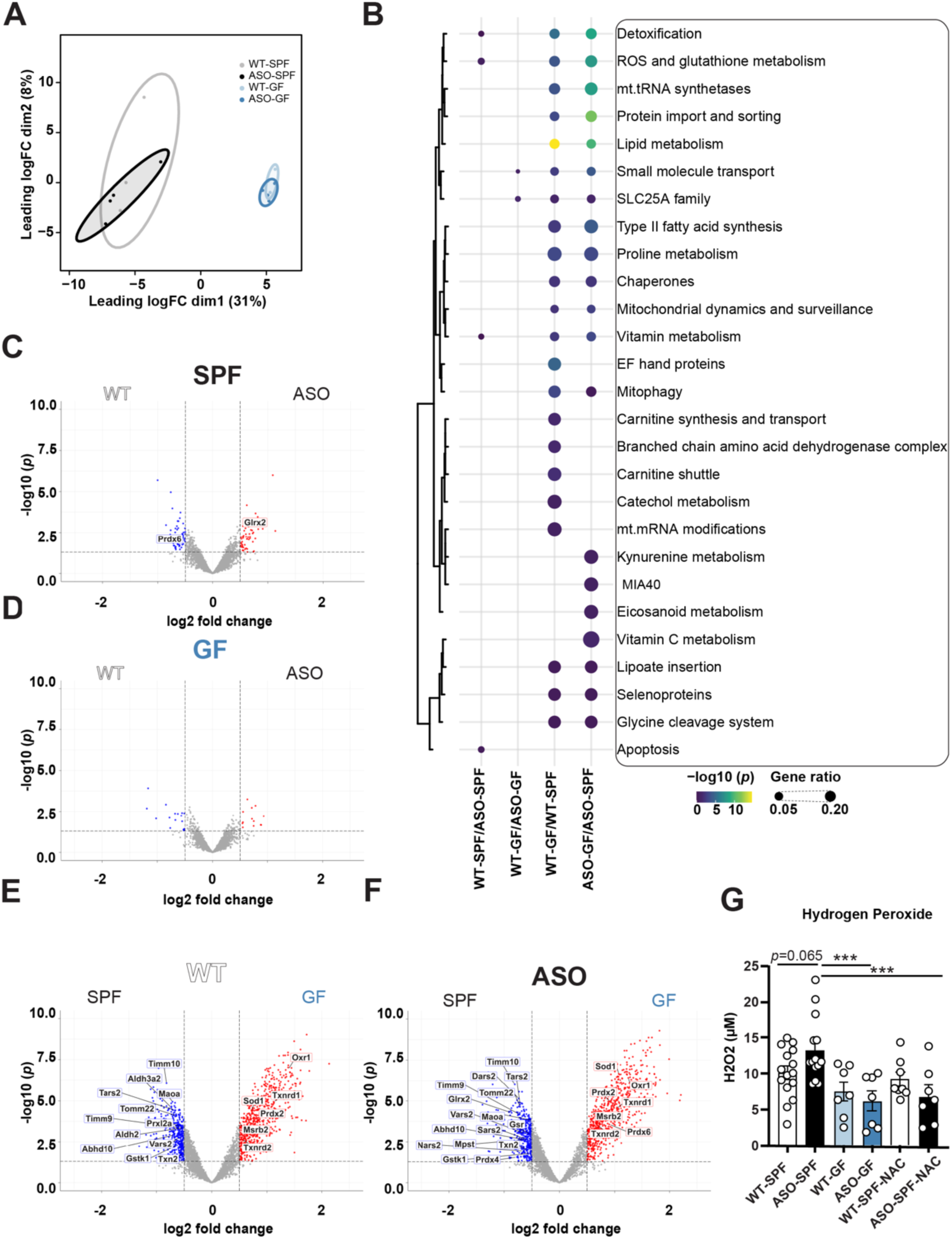
The gut microbiome regulates oxidative stress in ASO mice. **A-D)** Bulk proteomics of isolated mitochondrial extracts. (**A**) MDS plot of protein expression for different groups with 80% confidence intervals. (**B**) Top 10 most significantly altered pathways for each comparison, obtained through pathway analysis using the MitoCarta3.0 database. When groups were compared directly, each comparison produced 10 terms; however, upon consolidating all data, there were only 27 unique terms identified due to overlap between comparisons. (**C-F)** Volcano plots highlighting significantly altered proteins only from the top 5 differentially expressed pathways (detoxification, ROS and glutathione metabolism, mitochondrial tRNA synthetases, protein import and sorting, and lipid metabolism) based on (**C-D**) genotype and (**E-F)** microbiome. (**G**) H_2_O_2_ levels in the striatum measured using the cell-permeable fluorogenic probe DCFH-DA. SPF, specific pathogen-free; GF, germ-free; WT, wild-type; ASO, Thy1-α-synuclein overexpressing; NAC, N-Acetyl-cysteine; DCFH-DA, 2′-7′-dichlorodihydrofluorescein diacetate. **Statistical details:** (**A-F**) Proteins with *p*-value < 0.05 and |log_2_(fold change)| > 0.5 were identified as differentially expressed proteins. A two-sided *p*-value < 0.05 was used to determine statistically overrepresented pathways. Significance: *q* < 0.05. Sample size: n=4 mice per group. (**G**) Data were analyzed using a linear model (variable ∼ Genotype + Condition + Genotype*Condition) and pairwise comparisons with Benjamini-Hochberg (FDR) correction. Significance: *p* < 0.05. Data are expressed as mean ± SEM. Significance: *** *p* < 0.001. Sample size: n=7-15 mice per group.

Furthermore, mitochondrial tRNA synthetases such as DARS2 (aspartyl-tRNA synthetase 2, mitochondrial), TARS2 (threonyl-tRNA synthetase 2, mitochondrial), VARS2 (valyl-tRNA synthetase 2, mitochondrial), SARS2 (seryl-tRNA synthetase 2, mitochondrial), and NARS2 (asparaginyl-tRNA synthetase 2, mitochondrial) were enriched in mitochondria from the brains of SPF compared to GF animals, more so in ASO than WT mice (**Fig. 3E** and **F**).

Importantly, we discovered substantial enrichment of proteins involved in mitigating oxidative damage in germ-free mice (**Fig. 3E** and **F**). For example, proteins that control oxidative stress such as OXR1 (oxidation resistance 1), SOD1 (superoxide dismutase 1), FDPS (farnesyl diphosphate synthase), TXNRD1 (thioredoxin reductase 1), PRDX6 (peroxiredoxin 6), TXNRD2 (thioredoxin reductase 2), and ZADH2 (zinc binding alcohol dehydrogenase domain containing 2), among others, were significantly elevated in mitochondria from ASO-GF compared to ASO-SPF mice (**Fig. 3F**). We interpret this novel finding to suggest that GF αSyn-overexpressing mice maintain increased baseline expression of proteins to scavenge and neutralize ROS, potentially protecting animals from damage caused by altered mitochondrial function.

To determine whether the gut microbiome affects ROS levels, we measured H_2_O_2_ concentrations in the striatum using the cell-permeable fluorogenic probe DCFH-DA. Strikingly, significantly higher H_2_O_2_ levels were detected in ASO-SPF compared to ASO-GF mice (**Fig. 3G**). Treatment with the antioxidant N-acetylcysteine (NAC), a glutathione precursor, effectively reduced H_2_O_2_ levels in ASO-SPF mice to baseline levels (**Fig. 3G**). We conclude that the presence of a complex microbiome increases oxidative stress in the brain, potentially by reducing expression of antioxidant proteins in mitochondria, though we cannot exclude other mechanisms.

### Oxidative stress regulates motor function and Complex I activity in ASO-SPF mice

Altered levels of antioxidant molecules are of particular importance in PD progression (*69, 76, 77*), and elevated oxidative stress (i.e., lipid peroxidation) has been observed in ASO mice (*78*) though a link to the microbiome has not been previously reported. Accordingly, we found that neutralization of ROS via supplementation with NAC improves motor performance in ASO-SPF mice, as evidenced by decreased time to cross a challenge beam or to descend a pole, which is supported by mixture model analysis that shows NAC treated mice are more likely to successfully complete these motor behavior tasks (**Fig. 4, A and B**). Notably, antioxidant treatment had no impact on WT mice in either motor performance (**Fig. 4, A and B**) or striatal H_2_O_2_ levels (**Fig. 3G**).

**Figure 4.**
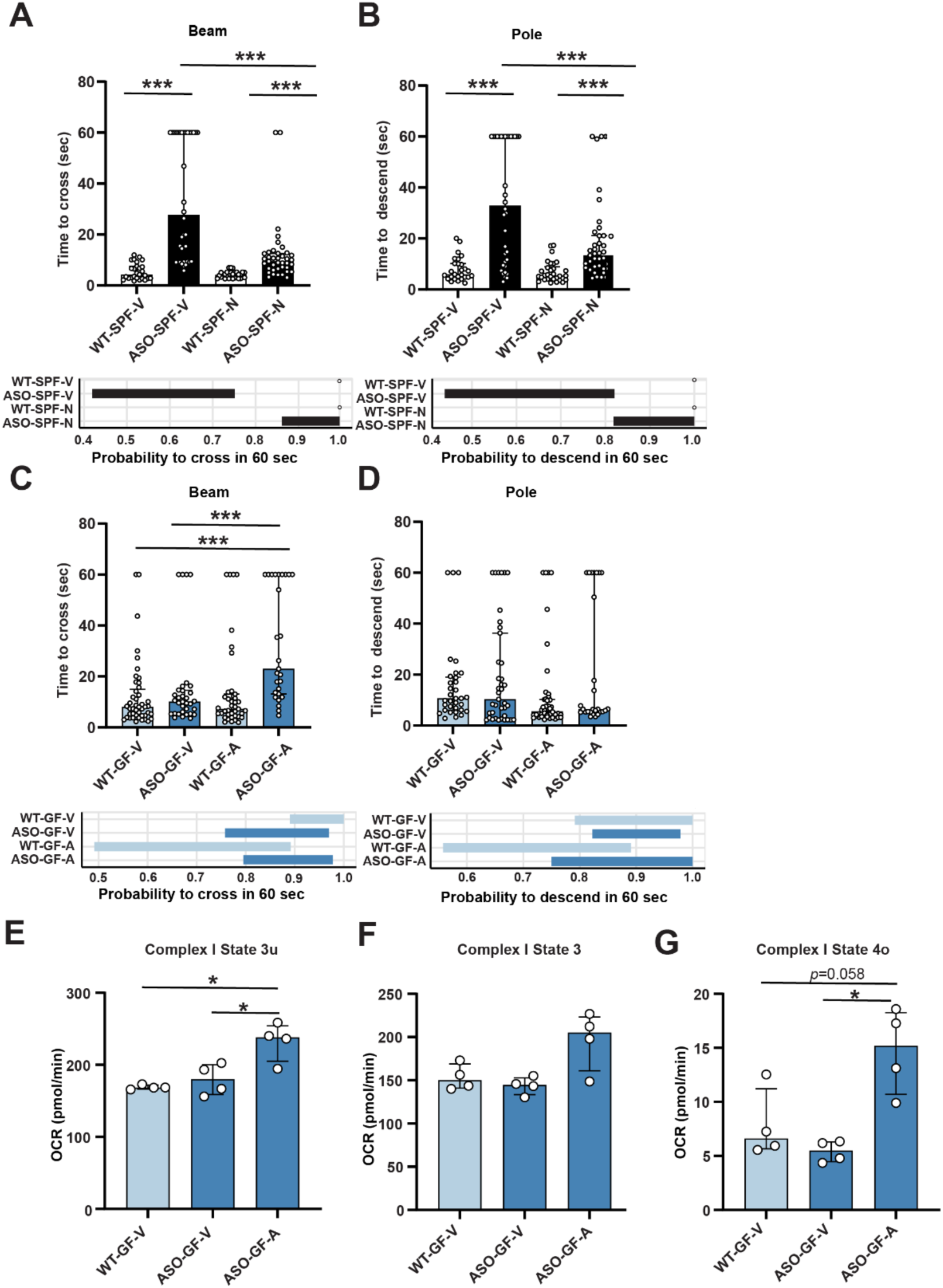
Oxidative stress regulates motor function and Complex I activity in ASO mice. (**A-D**) Treatment with N-Acetyl cysteine (NAC, abbreviated N) ameliorates motor behavioral symptoms in ASO-SPF mice, while inhibition of thioredoxin reductase with auranofin (abbreviated A) induces motor symptoms in ASO-GF mice. (**A, C**) Challenging beam test: time to cross (top) and 95% confidence interval (CI) of probability to cross successfully within 60 seconds (bottom) (**B, D)** Pole test: time to descend (top) and 95% CI of probability to descend successfully within 60 seconds (bottom). (**E-G**) Complex I oxygen consumption rate (OCR) measured in freshly isolated mitochondria using Seahorse. **E**) State 3u (uncoupled, maximal respiration). (**F**) State 3 (ATP-linked respiration). (**G**) State 4o (proton leak). SPF, specific pathogen-free; GF, germ-free; WT, wild-type; ASO, Thy1-α-synuclein overexpressing; NAC/ N, N-Acetyl-cysteine; V, vehicle; A, auranofin. **Statistical details:** (**A-D**) Bar plots show time to descend the pole the pole/cross the beam. Data are shown as median with interquartile range. Pairwise comparisons were calculated with Kruskal-Wallis test followed by Conover’s post-hoc test. Beam traversal and pole descent times were analyzed with a generative mixture model (y ∼ ωLogNormal(μ, σ) + (1 - ω)δ_y60_). CIs were calculated using maximum likelihood estimation (MLE) followed by parametric bootstrap. The ω 95% CI parameter describes the probability to descend the pole/cross the beam within 60 sec. Significance: *** *p* < 0.001. Sample size: n=9-13 mice per group, 3 trials per mouse. Data combined from 2 different cohorts. (**E-G**) Data were analyzed by Kruskal-Wallis test followed by Dunn’s post-hoc test with Benjamini-Hochberg (FDR) correction. Significance: * *p* < 0.05. Data are expressed as median ± interquartile range. Sample size: n= 4 mice per group.

Thioredoxin is one of the primary oxidation-reduction (redox) antioxidant systems in cells, along with glutathione. We observed that thioredoxin reductase (TXNRD)-1 and TXNRD-2 levels were upregulated in the mitochondrial proteome of ASO-GF mice (**Fig. 3F**), suggesting that elevated levels of ROS may overcome this antioxidant system and engender motor deficits.

Remarkably, experimental inhibition of TXNRD with auranofin in ASO-GF mice was sufficient to confer motor symptoms in the absence of a microbiota, significantly worsening performance in the challenge beam test (**Fig. 4C**). Further, WT animals treated with auranofin did not show motor deficits, confirming the requirement for αSyn overexpression to elicit PD-like symptoms (**Fig. 4C** and **D**). Interestingly, disruption of redox homeostasis by inhibition of thioredoxin reductase in ASO-GF mice selectively increased State 3u (uncoupled respiration) and State 4o (proton leak) in Complex I (**Figs. 4, E to G**, **S4, A and B**) without affecting Complex II oxygen consumption (**fig. S4C to G**). We thus uncover that GF mice exhibit an intrinsic ROS buffering activity that protects them from motor deficits induced by αSyn overexpression, and conclude that the microbiome contributes to PD-like pathophysiology via increasing oxidative stress in the ASO mouse brain.

## DISCUSSION

Evidence from multiple clinical cohorts across several geographies consistently reveal that gut microbiome composition in PD patients differs from that of healthy individuals (*36–40*). PD-associated microbiomes harbor increased levels of pro-inflammatory microbes and depletion of anti-inflammatory bacteria taxa (*37, 40, 79, 80*). Though most human studies can only predict putative functional outcomes, we previously reported that transplant of human microbiota from PD patients into ASO mice led to worse motor symptoms than transfer of intact gut microbial communities from non-PD human donors, suggesting microbiome changes may impact motor function (*48*). This finding was subsequently replicated in a toxicant-induced model (*81*). Several genetic risk factors for PD have been identified, including genes involved in mitochondrial biology and respiration, lysosomal biogenesis, autophagy, inflammation, and mutations in the gene encoding αSyn. However, many individual gene variants have low penetrance in the human population and only 15-20% of PD cases can be attributed to a monogenic cause. Therefore, most PD cases are idiopathic and likely arise from interactions between genetic and environmental risk factors such as pesticides, toxins, diet/lifestyle, and/or the microbiome (*82*).

We employed αSyn-overexpressing mice since aggregated forms of αSyn can disrupt mitochondrial biology through direct interaction with mitochondrial membranes, leading to altered mitochondrial dynamics, impairment of Complex I of the ETC, and increased oxidative stress via overproduction of ROS (*18–22*). Furthermore, αSyn aggregation can impair mitochondrial function in neurons, interfere with mitophagy, inhibit the autophagic degradation of damaged mitochondria, and contribute to cellular energy deficits and neuronal death (*20, 83*), particularly in dopaminergic neurons of the substantia nigra and striatum which are highly energy-dependent (*84*). Several additional interactions have been reported at the nexus of αSyn aggregation and mitochondrial function (*75, 85–87*).

To objectively explore interactions between αSyn and the gut microbiome, we profiled global transcriptomes in the striatum of ASO mice under SPF and GF conditions and discovered that gene expression signatures for respiratory electron transport, ATP synthesis, and the citric acid cycle were some of the most differentially expressed by gene-microbiome interactions. We experimentally validated this inference by directly showing increased respiration in ASO mice compared to WT animals or ASO mice devoid of a microbiome (i.e., germ-free ASO mice), with mitochondrial Complexes I and II most dramatically affected. We observed a similar increase in proton leak, indicating that part of the proton gradient from the ETC is being dissipated, which may reflect a protective cellular response against excessive ROS production. We did not detect changes in mitochondrial numbers, suggesting that at 4 months of age when ASO mice do not yet show neurodegeneration (*55*), individual mitochondria in the striatum are overactive due to the burden of αSyn accumulation. We speculate that this phenotype is a compensatory mechanism at this early age, when cells are stressed due to aggregation of αSyn and respond by increasing respiration to meet their energy demands. Indeed, defects in OxPhos can drive mitochondrial hypermetabolism, triggering stress responses and reducing cell viability (*73, 88*). Our findings converge with evidence that mitochondrial hyperactivity can precede neuronal loss and represents an early pathological event in PD (*89*), though further studies are needed to conclude how increased mitochondrial respiration and neurodegeneration are functionally linked.

αSyn-driven mitochondrial stress persisting for months to years may result in neuronal dysfunction and ultimately cell death/neurodegeneration. Of note, we do not know which cell types are most affected by mitochondrial hyperfunction in ASO mice—the neurons that overexpress αSyn and/or adjacent cells that respond to neuronal stress.

Untargeted proteomic analysis of isolated mitochondria revealed several processes dysregulated in ASO mice, in a microbiome-dependent manner, including lipid metabolism, protein import, mitochondrial tRNA synthetases, detoxification pathways, and ROS and glutathione metabolism. Mitochondria are a significant source of ROS, which include byproducts of the ETC during oxidative phosphorylation such as superoxide anions (O_2_^•-^), H_2_O_2_, and hydroxyl radicals (^•^OH) (*90*). Excessive ROS production promotes oxidative stress, damaging cellular components such as lipids, proteins, and DNA (*77*). Impaired ETC function and increased electron leakage exacerbate ROS production, which can in turn further impair mitochondrial function (*90*). Elevated levels of ROS have been broadly implicated in PD, associated with mitochondrial dysfunction caused by genetic mutations (*PINK1, PARK2*, and *PARK7*) or toxins such as rotenone or 1-methyl-4-phenyl-1,2,3,6-tetrahydropyridine (*77*). Indeed, we show here for the first time that interaction between αSyn overexpression and the gut microbiome promotes increased H_2_O_2_ in the brain, and that ROS production is critical for PD-related outcomes, as treatment with the antioxidant NAC improves motor symptoms in ASO mice.

Mitochondria have several antioxidant defense mechanisms to mitigate ROS damage, including enzymes such as superoxide dismutase (SOD), catalase, glutathione peroxidase, and thioredoxins and TXNRD (*91*). Strikingly, we found that many of these antioxidant proteins are upregulated in striatal mitochondria harvested from GF mice, of both WT and ASO genotypes, suggesting an intrinsic default process to protect cells from oxidative damage that can be overwhelmed by microbial colonization. In other words, the gut microbiome promotes αSyn aggregation in ASO mice (*59*) and in turn mitochondrial dysfunction, leading to elevated ROS levels. The epistasis of these events requires further study. Importantly, treatment of ASO-GF mice with a TXNRD inhibitor to experimentally block antioxidant defense capacity in uncolonized ASO mice induced motor phenotypes, with a concomitant increase in Complex I respiration in the striatum. Thus, of the many potential contributions by a complex gut microbiome, elevating ROS in the brain appears sufficient to induce motor symptoms in mice that overexpress αSyn. If validated in other animal models and in humans, microbiome modulation of oxidative stress may be an environmental risk underlying the pathophysiology of PD. Further, microbiome-mediated production of ROS uncovers targets for novel PD interventions that drug the gut, rather than the more challenging prospect of therapeutics directed to the brain.

## MATERIALS AND METHODS

### Mice

Male mice overexpressing human αSyn under the Thy1 promoter (“Line 61” Thy1-αSyn, ASO) and WT mice were reared by crossing WT BDF1 males (Charles River, RRID:IMSR_CRL:099) with ASO heterozygous females (*55*). Since the transgene is carried on the X chromosome, only male animals were used to avoid the effects of X-inactivation. GF mice were born by caesarean section, with the offspring fostered by GF Swiss-Webster dams and kept microbiologically sterile inside flexible film isolators. SPF mice were housed in autoclaved micro-isolator cages. GF status was verified on a bi-weekly basis through 16S rRNA PCR of fecal-derived DNA and plating of fecal pellets on Brucella blood agar under anaerobic conditions and tryptic soy blood agar under aerobic conditions. All mice received food and water *ad libitum*, were maintained on the same 12-hour light-dark cycle, and were kept in the same animal facility. All animal husbandry and experiments were authorized by the California Institute of Technology’s Institutional Animal Care and Use Committee (IACUC). All experiments were performed when the mice were 4 months of age.

### Pharmacological treatments

#### N-Acetyl-L-cysteine (NAC)

Mice were given ad libitum access to drinking water supplemented with 40 mM N-Acetyl-L-cysteine (NAC, N) (Sigma-Aldrich, Cat. #A7250) or water without treatment (vehicle, V) for 8 weeks, beginning at two months of age. Fresh water and NAC treatment were provided weekly.

#### Auranofin

Mice were administered Auranofin (Cayman, Cat. #15316), suspended in a 5% dimethyl sulfoxide (DMSO, 99.9%, diluted to 5% in saline) solution as the vehicle (V). The dose was 1 mg/kg, administered via intraperitoneal injection once daily for 5 consecutive days. A separate group of mice received only the vehicle solution.

### Motor testing

Experiments involving GF and GF x SPF comparisons were conducted inside isolators. For NAC intervention, experiments were completed in a biological safety cabinet. All experiments were performed between ZT6 and 10 of the light phase. Motor testing was performed as previously described (*92, 93*) and in the following order: Day 1: beam traversal training and pole training; Day 2: beam traversal training and pole training; Day 3: beam traversal test and pole test.

#### Pole descent

This test measures the time for mice to descend a 24-inch pole wrapped in mesh. Mice were trained for two days with three trials: 1) placed head down ⅓ of the height from the base, 2) placed head down ⅔ of the height from the base, and 3) placed head down at the top. On testing day, mice were placed at the top of the pole for three trials and the time to descend the pole was recorded. The time was recorded when the hindlimbs reached the base, with a maximum time of 60 seconds. Mice that fell or slid were given a score of 60 seconds. For protocol, see dx.doi.org/10.17504/protocols.io.n92ld8j87v5b/v1.

#### Challenging beam

This test measures the time for mice to cross a 1-meter plexiglass beam with segments decreasing in width (3.5 cm, 2.5 cm, 1.5 cm, 0.5 cm). Mice were trained for two days before testing. On testing day, mice were placed at the start of the 3.5 cm segment for three trials and the time to cross the beam was recorded. Time was recorded when hindlimbs reached the home cage, with a maximum time of 60 seconds. Mice that fell were given a score of 60 seconds. For protocol, see dx.doi.org/10.17504/protocols.io.e6nvw1212lmk/v1.

### Striatal tissue dissection

Mice were sacrificed by cervical dislocation. The brain was rapidly removed and placed in an ice-chilled stainless steel coronal matrix. Brain tissue was sectioned using a 1.0 mm brain matrix (Roboz, SA-2175). The striatum was dissected within three minutes using reference brain atlas coordinates. Coordinates for striatal brain slices spanned from anterior to posterior (AP) +1.54 mm to +0.10 mm relative to bregma. For protocol, see dx.doi.org/10.17504/protocols.io.x54v92z2ql3e/v1.

### Oxidative stress quantification

Following dissection, whole striatal tissue was homogenized in 100 µL PBS. H_2_O_2_ was quantified using the OxiSelect™ In Vitro ROS/RNS Assay Kit (Green Fluorescence) from CellBio Labs (Cat. #STA-347-5) according to the manufacturer’s instructions. Briefly, 50 µL of the sample (1:25 dilution in PBS) or H_2_O_2_ standard was added to a black 96-well plate in triplicate. Then, 50 µL of catalyst was added to each well, mixed, and the plate was incubated at room temperature for 5 minutes before 100 µL of DCFH solution was added to each well. The plate was protected from light and incubated at room temperature for 30 minutes. The fluorescence was read at 480 nm excitation/530 nm emission. For protocol, see dx.doi.org/10.17504/protocols.io.4r3l2qbqjl1y/v1.

### Enzymatic activity profiling

Striatal tissue samples were dissected, weighed and homogenized 1:180 (weight:volume, mg:µL) in homogenization buffer (50 mM Triethanolamine and l mM EDTA) with 2 tungsten beads (Qiagen, Cat. #69997) using a Tissue Lyser (Qiagen Cat. # 85300) at 30 cycles/sec for 1 minute, incubated on ice for 5 minutes, and re-homogenized for 1 minute. Homogenates were vortexed before each use to ensure homogeneity. Enzymatic activities were measured spectrophotometrically for Complex I (NADH-ubiquinone oxidoreductase), Complex II (succinate-ubiquinone oxidoreductase), Complex IV (cytochrome c oxidase) and CS (citrate synthase) as previously described (*94*), with minor modifications. All enzymatic assays were performed using l0 µL of brain homogenate in 96-well plates in a Spectramax M2 plate reader (Spectramax Pro 6, Molecular Devices). Samples were run in duplicate for each enzymatic activity assay, along with a non-specific activity control, and final enzymatic activities were determined by averaging the duplicates. The specific activity of each sample was calculated as the total activity minus the non-specific activity (negative control). To maximize the accuracy of comparisons between animals, all samples were run on the same plate, and five reference samples were used in each plate to adjust for batch/plate effects, using a derived normalization factor for each assay.

The mitochondrial health index (MHI) integrates 5 primary features, yielding an overall per-mitochondrion score of mitochondrial respiratory chain activity (*95*). The equation uses the mean-centered activity of Complexes I, II, and IV as a numerator, divided by two indirect markers of mitochondrial content, mtDNA density and CS activity: MHI = (CI+CII+CIV) / (CS+mtDNA density+1) 100. A value of 1 is added as a third factor in the denominator to balance the equation. Values for each of the 5 features are mean-centered (giving the value for an animal relative to all other animals) such that an animal with average activity for all features will have an MHI of 100 ([1+1+1] / [1+1+1] 100 = 100). The mitochondrial respiratory capacity (MRC) is a modified version of the MHI which takes into consideration the volumetric scaling of mitochondrial content metrics and the surface area scaling of cristae-bound ETC components.

MRC is computed from the cube root (for volume) and square root (for surface area) of the mitochondrial mass and respiratory chain, respectively, as (*71*). For protocol, see: dx.doi.org/10.17504/protocols.io.kxygxy2nwl8j/v1.

### Mitochondrial DNA quantification

mtDNA density and mtDNA copy number (mtDNAcn) were measured from the total homogenates used for the enzymatic activity measures. Brain homogenates were lysed in a Thermocycler at a 1:10 dilution in lysis buffer (114 mM Tris HCI pH 8.5, 6% Tween 20, and 200 μg/ml proteinase K) for 16 hours at 55°C, followed by heat inactivation for 10 minutes at 95°C, and were kept at 4°C until used for qPCR, as described previously with minor modifications (*94*). qPCR reactions were performed in triplicate in 384-well qPCR plates with 12 μl of master mix (TaqMan Fast Universal Master Mix, Life Technologies Cat. #4444964) and 5 μl of lysate. Mitochondrial and nuclear amplicons were quantified in the same reaction: cytochrome c oxidase subunit 1 (COXl, mtDNA) and beta-2 microglobulin (B2M, nDNA). The Master Mix included 300 nM of primers and 100 nM of probe: COXl-Fwd: ACCACCATCATTTCTCCTTCTC, COXl-Rev: CTCCTGCATGGGCTAGATTT, COXl-Probe: HEX/AAGCAGGAG/ZEN/CAGGAACAGGATGAA/3IABkFQ. B2M-Fwd: GAGAATGGGAAGCCGAACATA, B2M-Rev: CCGTTCTTCAGCATTTGGATTT, B2M-Probe: FAM/CGTAACACA/ZEN/GTTCCACCCGCCTC/3IABkFQ. The mtDNAcn was derived from the Λ1 Cycle threshold (Ct), calculated by subtracting the average mtDNA Ct from the average nDNA Ct. mtDNAcn was calculated by 2^(βCt^) x 2 to account for the diploid nuclear genome. Measures of mtDNA density were derived by linearizing the Ct values as 2^Ct^ / (1/10^-12^) to give relative mtDNA abundance per unit of tissue. For protocol, please see: dx.doi.org/10.17504/protocols.io.14egn6jwzl5d/v1.

### Isolated mitochondrial activity profiling

#### Isolation

Striatal tissues were placed in 1 mL ice-cooled MSHE buffer (210 mM mannitol, 70 mM sucrose, 5 mM HEPES, 1 mM EGTA, 0.2% fatty acid-free BSA) immediately after dissection, then transferred to a 15 mL Dounce homogenizer and homogenized with ten pestle strokes. The homogenate was centrifuged at 2,000×g for 3 minutes at 4°C. The supernatant was collected, placed in a new tube, and centrifuged at 12,000×g for 10 minutes at 4°C. The supernatant was aspirated, and the pellet was resuspended in 10% digitonin plus MSHE before being centrifuged again at 12,000×g for 10 minutes at 4°C. After removing the supernatant and upper white layer, the pellet was resuspended in MSHE without BSA and centrifuged once more. Finally, all the supernatant and remaining white layer above the pellet were removed, and the mitochondria-enriched pellet was resuspended in MSHE without BSA for experiments. For protocol, see dx.doi.org/10.17504/protocols.io.5jyl82qz6l2w/v1.

#### Respirometry

Three to 5 μg of mitochondrial extracts were plated at 20 µL per well. The plate was then centrifuged at 2,100×g for 5 minutes at 4°C (without brake), followed by the addition of 130 μL of MAS (containing 220 mM mannitol, 70 mM sucrose, 5 mM KH_2_PO_4_, 5 mM MgCl_2_, 2 mM HEPES, 1 mM EGTA, and 0.1% (w/v) fatty acid-free BSA) + substrate to each well. The following substrates were used to stimulate mitochondrial respiration: 5 mM pyruvate and 1 mM malate (for Complex I respiration), and 5 mM succinate with 2 µM rotenone (for Complex II respiration). To assess various bioenergetic parameters, the following compounds were injected: 3 μM ADP, 5 μM oligomycin, 4 μM FCCP, and 2 μM Rotenone + 2 μM Antimycin A. The oxygen consumption rates (OCR) were measured using the Seahorse XF96 extracellular flux analyzer (Agilent Technologies, Santa Clara, CA). Measurements were performed in quintuplicate and averaged for each mouse. The obtained OCR rates were normalized by protein concentration. For protocol, see dx.doi.org/10.17504/protocols.io.bp2l62z4zgqe/v1.

### Western blotting

Whole striatal tissue and mitochondrial extracts were kept at −80°C until use. Whole striatal tissue was homogenized in 1X RIPA buffer plus protease and phosphatase inhibitors with a hand tissue homogenizer (BT704). Protein levels were determined using Western blot analysis. Protein concentration for each sample was determined using the Pierce BCA Protein Assay Kit (ThermoFisher) and normalized to 1 µg/µL in PBS. Twenty micrograms of protein were separated by SDS-PAGE using 4-20% Tris-Glycine WedgeWell gels and transferred onto a 0.45 µm PVDF membrane. Beta actin was used as a loading control. Blocking was performed in 5% non-fat dry milk in Tris-buffered saline with 0.1% Tween-20, followed by overnight staining at 4°C with primary antibodies recombinant anti-TOMM20 (Abcam, #ab186735), anti-Grp75 (Abcam, # ab2799), anti-mtTFA (Abcam, # ab 252432), total OXPHOS Rodent WB Antibody Cocktail (Abcam, #ab110413), and anti-VDAC1 (Abcam, # #15895) at a dilution of 1:1000. The next day, the blots were stained with secondary antibodies anti-rabbit IgG, HRP-linked (Cell signaling, # 7074), and anti-mouse IgG, HRP-linked (Cell signaling, #7076) (1:1000) for 1.5 hours, and the signal was detected with Clarity chemiluminescence substrate before imaging on a Bio-Rad imager. For protocol, see dx.doi.org/10.17504/protocols.io.x54v92z2ql3e/v1.

### Quantitative bulk transcriptomics

#### Sequencing

RNA sequencing was performed on total striatal tissue extracts. After dissection, striatum was stored in RNAlater (Qiagen) at 4°C for 24 hours, then RNAlater was removed, and samples were snap-frozen in dry ice and stored at −80°C until RNA isolation. Striatal tissue was homogenized in TRIzol (Ambion) with a pestle. Total RNA was isolated by adding chloroform to the samples and centrifuging at 12,000×g for 15 minutes. The aqueous layer was further purified using the RNeasy Mini Kit (Qiagen) following the manufacturer’s instructions. RNA integrity numbers (RINs) were assessed with a Bioanalyzer (Agilent), and samples with low RINs were not sequenced. RNA-seq libraries were prepared in the Penn State College of Medicine Genome Sciences core (RRID:SCR_021123) using Lexogen’s QuantSeq 3’ mRNA-Seq Library Prep Kit Forward (FWD) following the manufacturer’s instructions. Briefly, total mRNA was reverse transcribed using oligo (dT) primers. The second cDNA strand was synthesized by random priming followed by cDNA purification and library amplification using single indices. The libraries were analyzed for size distribution and concentration using the BioAnalyzer High Sensitivity DNA kit (Agilent technologies, Santa Clara, CA). Libraries were pooled at equimolar concentrations and sequenced on a Novaseq6000 (Illumina) to get approximately 12 million, single end reads. Read quality was validated with FastQC (0.11.6, http://www.bioinformatics.babraham.ac.uk/projects/fastqc/, RRID:SCR_014583). Phred scores for reads were greater than 20, and no GC bias was detected in any samples. For protocol, see dx.doi.org/10.17504/protocols.io.3byl4w7rzvo5/v1.

#### Data analysis

A custom genome for the ASO mouse was generated by splicing human αSyn (nt 53-475; hSNCA) into the mouse genome (GRCm39, Ensembl release 110). Rsubread package v2.16.1 in (https://bioconductor.org/packages/release/bioc/html/Rsubread.html, RRID:SCR_016945) was used in R (v4.3.2) to align trimmed reads against this custom genome and *featureCounts* (https://subread.sourceforge.net/featureCounts.html, RRID:SCR_012919) used with default settings to generate gene level counts (*96*). Enrichment for different cell types was assessed by gene expression deconvolution using dampened weighted least squares with the DWLS package package v0.1.0 (https://github.com/dtsoucas/DWLS)(97). Striatal cells from Zhang et al. (2023) (*98*) were used as a reference for deconvolution. Data were normalized with DESeq2 v1.42.1 (https://bioconductor.org/packages/release/bioc/html/DESeq2.html, RRID:SCR_015687)(*99*) and principal coordinate analysis was subsequently performed by multidimensional scaling to assess sample level clustering using the limma package (v3.60.4, RRID:SCR_010943)(*100*). DESeq2 was then used to assess differential gene expression between (**1**) WT and ASO mice separately in the SPF and GF conditions (assessing effect of genotype) and (**2**) SPF and GF conditions separately in WT and ASO mice (assessing effect of the microbiome). A two-tailed false discovery rate (*FDR*) *< 0.05* (original Benjamini-Hochberg method) determined statistical significance and |log_2_(FC)*| > 0.5* was selected to identify differentially expressed genes (DEGs). Gene set enrichment analysis (GSEA) was performed with DEGs separately for upregulated and downregulated genes for each comparison against the Kyoto Encyclopedia of Genes and Genomes (KEGG), Gene Ontology (GO) and Reactome pathways using the Rapid Integration of Term Annotation and Network (RITAN) package v1.26.0 (https://bioconductor.org/packages/release/bioc/html/RITAN.html)(101). A two-sided adjusted p-value (q-value) threshold of 0.05 (Benjamini-Hochberg method) was used to determine statistically overrepresented pathways.

### Proteomics

#### Sample preparation

After mitochondrial isolation, 100 µg of protein derived from mitochondrial-enriched fractions were digested with S-trap™ (Protifi, Fairport, NY) according to the manufacturer’s protocol. Briefly, the protein was reduced with tris(2-carboxyethyl)phosphine, alkylated with chloroacetamide, and digested with trypsin overnight. The resulting peptides were eluted from the S-trap according to the manufacturer’s protocol. The eluates were dried and resuspended in 2% acetonitrile, 0.2% formic acid in water. For protocol, see dx.doi.org/10.17504/protocols.io.36wgqny15gk5/v1

#### LC-MS/MS

LC-MS/MS experiments were performed by loading 500 ng of sample onto an EASY-nLC 1200 (ThermoFisher Scientific, San Jose, CA) connected to a Q Exactive HF Quadrupole-Orbitrap Hybrid mass spectrometer (Thermo Fisher Scientific, San Jose, CA). Peptides were separated on an Aurora UHPLC Column (25 cm × 75 µm, 1.6 µm C18, AUR2-25075C18A, Ion Opticks) with a flow rate of 0.35 µL/min for a total duration of 131 minutes. The gradient was composed of 3% Solvent B for 1 minute, 3–19% B for 72 minutes, 19–29% B for 28 minutes, 29–41% B for 20 minutes, 41–95% B for 3 minutes, and 95–98% B for 7 minutes. Solvent A consisted of 97.8% H_2_O, 2% ACN, and 0.2% formic acid, and solvent B consisted of 19.8% H_2_O, 80% ACN, and 0.2% formic acid. MS1 scans were acquired with a range of 375–1500 m/z in the Orbitrap at 60,000 resolution. The maximum injection time was 15 ms, and the AGC target was 3 × 10^6^.

MS2 scans were acquired at 30,000 resolution with a first scan mass of 100 Da. The maximum injection time was 45 ms, and the AGC target was 3 × 10^6^. The isolation window was 1.2 m/z, collision energy was 28 NCE, and loop count was 12. Other global settings were as follows: ion source type, NSI; spray voltage, 2000 V; ion transfer tube temperature, 300°C. Method modification and data collection were performed using Xcalibur software (Thermo Scientific, http://chemistry.unt.edu/~verbeck/LIMS/Manuals/XCAL_Quant.pdf, RRID:SCR_014593). For protocol, see dx.doi.org/10.17504/protocols.io.36wgqny15gk5/v1

#### Data analysis

Data analysis was performed using Proteome Discoverer (v2.5, Thermo Scientific, https://www.thermofisher.com/order/catalog/product/IQLAAEGABSFAKJMAUH, RRID:SCR_014477) software, the Uniprot mouse database from UniProtKB (http://www.uniprot.org/help/uniprotkb, RRID:SCR_004426), and Sequest with Percolator (http://noble.gs.washington.edu/proj/percolator/, RRID:SCR_005040) validation. Normalization was performed with random-forest normalization using the tidyproteomics R package (https://jeffsocal.github.io/tidyproteomics/). Principal coordinate analysis was performed by multidimensional scaling to assess sample level clustering using the limma package (v3.60.4, RRID:SCR_010943)(*98*). Differentially expressed proteins were identified using a two-sided Student’s t-test implemented in the limma package (v3.60.4). Proteins with a *p-value < 0.05* were considered statistically significant, and those with *|log2(fold change)| > 0.5* were identified as differentially expressed proteins. Pathway analysis was performed for each comparison against the mouse MitoCarta database (3.0, http://www.broadinstitute.org/pubs/MitoCarta/, RRID:SCR_018165) using the RITAN package. MitoCarta inventories and annotates genes with mitochondrial localization, and was thus used to more precisely identify the involvement of specific mitochondrial pathways. A two-sided *p*-value < 0.05 was used to determine statistically overrepresented pathways.

### Statistical analysis

#### Motor testing

Data analysis was conducted in Python (v3.12.6, http://www.python.org, RRID:SCR_008394) with the following packages: NumPy (v2.0.2, http://www.numpy.org, RRID:SCR_008633), Pandas (v2.2.3, https://pandas.pydata.org, RRID:SCR_018214), Bokeh (v3.5.2, https://bokeh.org), bebi103 (v0.1.25, https://bebi103.github.io) and JupyterLab (v4.2.5, http://jupyterlab.github.io/jupyterlab/, RRID:SCR_023339). The full list of required packages with versions can be found in a .yml file in the GitHub repository associated with this paper. Data from all trials were considered independent and identically distributed and hence were pooled for analysis. A score of 60 s was assigned to all the mice that did not finish the test in 60 s. Analysis was performed with custom Python code. The time to descend the pole or cross the beam was fit into a right-censored lognormal mixture model:

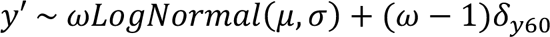

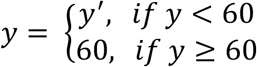

where ω (*omega*) is a parameter describing the probability of a subject mouse finishing the test successfully (descending the pole or crossing the beam); μ (*mu*), σ(*sigma*) are standard parameters of the LogNormal distribution; δ_y60_ is the Kronecker delta function. The model’s fit was assessed through a graphical model assessment approach (Q-Q plots and predictive ECDFs). The parameters of the model were estimated using the maximum likelihood method. Parameter maximum likelihood estimates (MLEs) were then used to generate 10,000 bootstrap replicates.

Parameter confidence intervals were built for group comparisons. *p*-values were calculated using the Kruskal-Wallis test followed by the post-hoc Conover’s test.

#### Bioenergetics and biochemical assays

Data analysis was conducted using R (v4.3.2, https://www.r-project.org/, RRID:SCR_001905). For respirometry, oxidative stress quantification, mitochondrial enzymatic activities, mitochondrial DNA quantification, and Mitochondrial Health Index, we used a linear model *(variable ∼ Genotype + Microbiome + Genotype*Microbiome)* and conducted pairwise comparisons with Benjamini-Hochberg (FDR) correction. For immunofluorescence data and other datasets with sample sizes smaller than six, we performed a Kruskal-Wallis test followed by Dunn’s post-hoc test with Benjamini-Hochberg (FDR) correction. For H_2_O_2_ assay data we used a linear model *(variable ∼ Genotype + Condition + Genotype*Condition)* and conducted pairwise comparisons with Benjamini-Hochberg (FDR) correction.

## Supporting information

Source Data for Figure 3A-F

## ACKNOWLEDGMENTS

We thank Taren Thron for assistance with animal experiments and animal breeding, Yvette Garcia-Flores for logistical assistance, Dr. Catherine Oikonomou for comprehensive editing of the manuscript, and the Mazmanian laboratory for critical feedback on the research and manuscript.

## Author contributions

Conceptualization: LHM, SKM

Methodology: LHM, LS, ADO, JDH, JSB, T-FC, JT, MP

Investigation: LHM, LS, MF, ADO, JDH, JJ, BQ, JD

Visualization: LHM, ADO, JDH

Supervision: T-FC, JT, MP, VG, SKM

Writing–original draft: LHM, SKM

Writing–review & editing: LHM, LS, ADO, JDH, JD, JSB, T-FC, JT, MP, VG, SKM

## Data and materials availability

The data, code, protocols, and key lab materials used and generated in this study are listed in a Key Resource Table alongside their persistent identifiers (Table S2). Tabular data and analysis code are available at https://github.com/jdthoang/LHMorais2024. The brain transcriptomics dataset has been deposited and is accessible via BioProject accession number PRJNA1181029 in the NCBI BioProject database (https://www.ncbi.nlm.nih.gov/bioproject/). The proteomics dataset is available from doi:10.25345/C53B5WM0M with the MassIVE identifier MSV000096259 and the ProteomeXchange identifier PXD057375.

