## Supplementary Information for "The gut microbiome promotes mitochondrial respiration in the brain of a Parkinson’s disease mouse model"

Livia H. Morais *et al.*

**This PDF file includes:**

Figs. S1 to S4
Table S2

**Other Supplementary Materials for this manuscript include the following:**

Table S1: Differentially expressed genes between groups
Estimated log2 fold changes and p-values for genes differentially expressed in various group comparisons.
Source data for Figure 3A-F

FIGURE S1

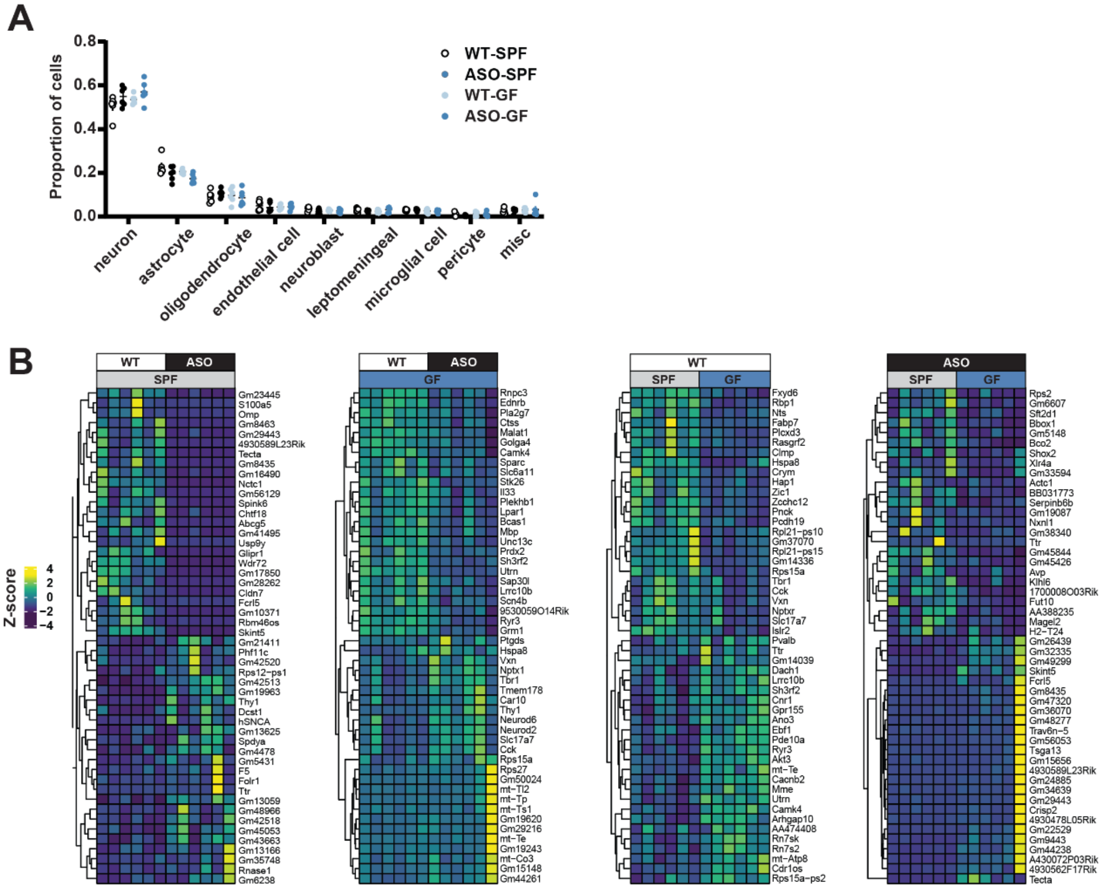


**Fig. S1. No differences in cell-type enrichment across groups.**

Bulk quantitative RNA-sequencing of whole striatum tissue. Differential gene expression between groups was assessed using the DESeq2 package comparing the effects of genotype (WT x ASO) and the gut microbiome (SPF x GF). (**A**) Cell-type enrichment analysis of all differentially expressed genes (DEGs) detected no differences between groups. (**B**) Heat map showing the top 25 up- and downregulated genes in each comparison. SPF, specific pathogen-free; GF, germ-free; WT, wild-type; ASO, Thy1-α-synuclein overexpressing.

FIGURE S2

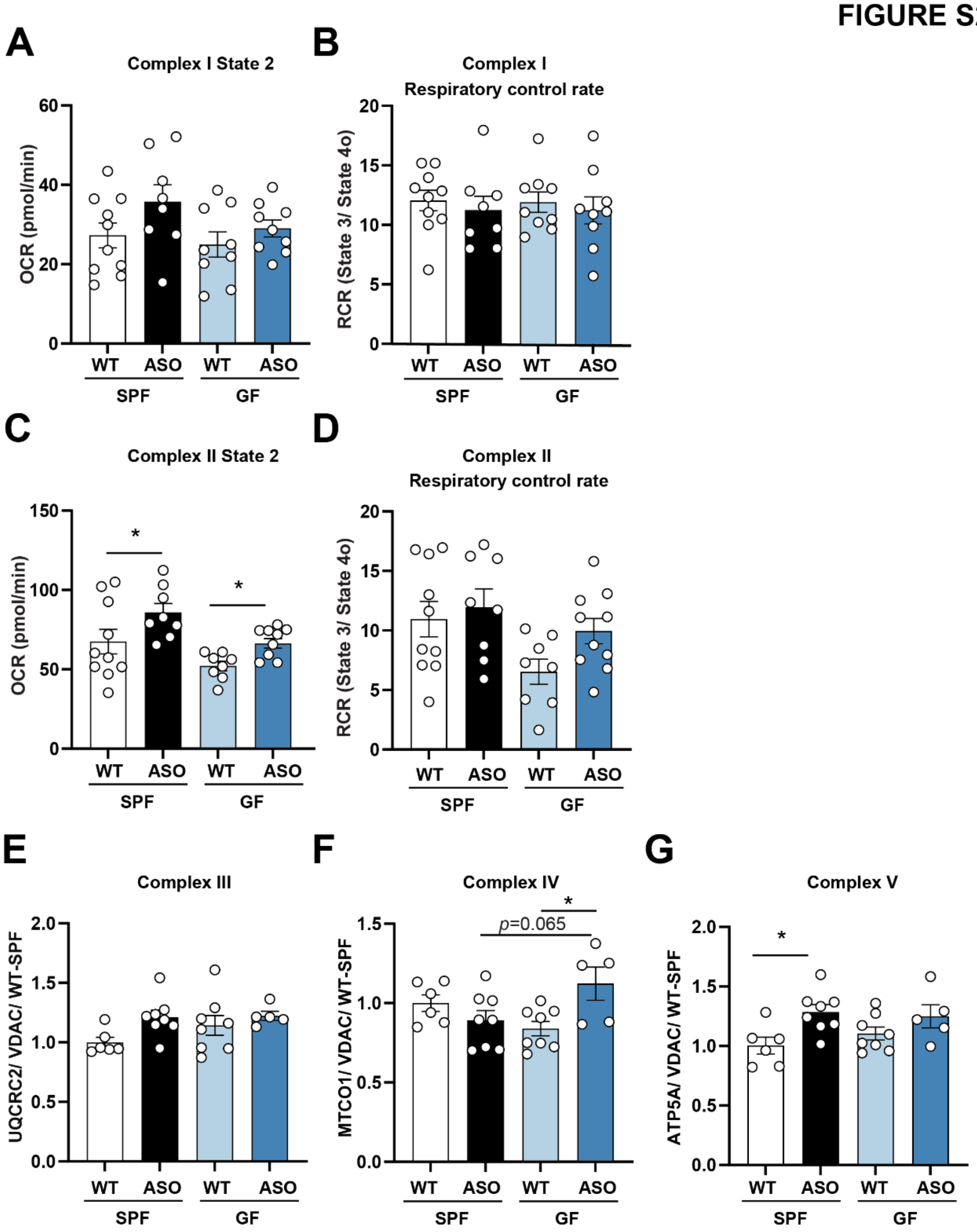
A
Complex I State 2
OCR (pmol/min)
WT
ASO
SPF
GF
B
Complex I
Respiratory control rate
RCR (State 3/ State 4o)
C
Complex II State 2
D
Complex II
Respiratory control rate
E
Complex III
UQCRC2/ VDAC/ WT-SPF
F
Complex IV
p=0.065
MTCO1/ VDAC/ WT-SPF
G
Complex V
ATP5A/ VDAC/ WT-SPF

**Fig. S2. Gut microbiome regulates mitochondrial respiration in ASO mice.**

(**A-D**) Oxygen consumption rate (OCR) was measured in freshly isolated striatal mitochondria using Seahorse, evaluating Complex I and Complex II respiration. (**A, C**) State 2. (**B, D**) Respiratory control rate (RCR). (**E-G**) Quantification of levels of oxidative phosphorylation (OxPhos) complexes, normalized to voltage-dependent anion channel 1 (VDAC1), in isolated mitochondria by Western blotting. (**E**) Complex III UQCRC2 subunit. (**F**) Complex IV MTCO1 subunit. (**G**) Complex V ATP5A subunit. SPF, specific pathogen-free; GF, germ-free; WT, wild-type; ASO, Thy1-α-synuclein overexpressing; RCR, respiratory control rate; CI, Complex I; CII, Complex II; CIII, Complex III; CIV, Complex IV; CV, Complex V; UQCRC2, ubiquinol-cytochrome c reductase core protein 2; MTCO1, cytochrome c oxidase subunit 1; ATP5A, ATP synthase subunit alpha.

**Statistical details**: (**A-G**) Data were analyzed using a linear model (variable ~ Genotype + Microbiome + Genotype*Microbiome) and pairwise comparisons with Benjamini-Hochberg (FDR) correction. Data are expressed as mean ± SEM. Significance: * $p < 0.05$. (**A-D**) Sample size: n=8-10 per group, data combined from 4 different cohorts. (**E-G**) Sample size: n=6-8 per group, data combined from 2 different cohorts.

### FIGURE S3

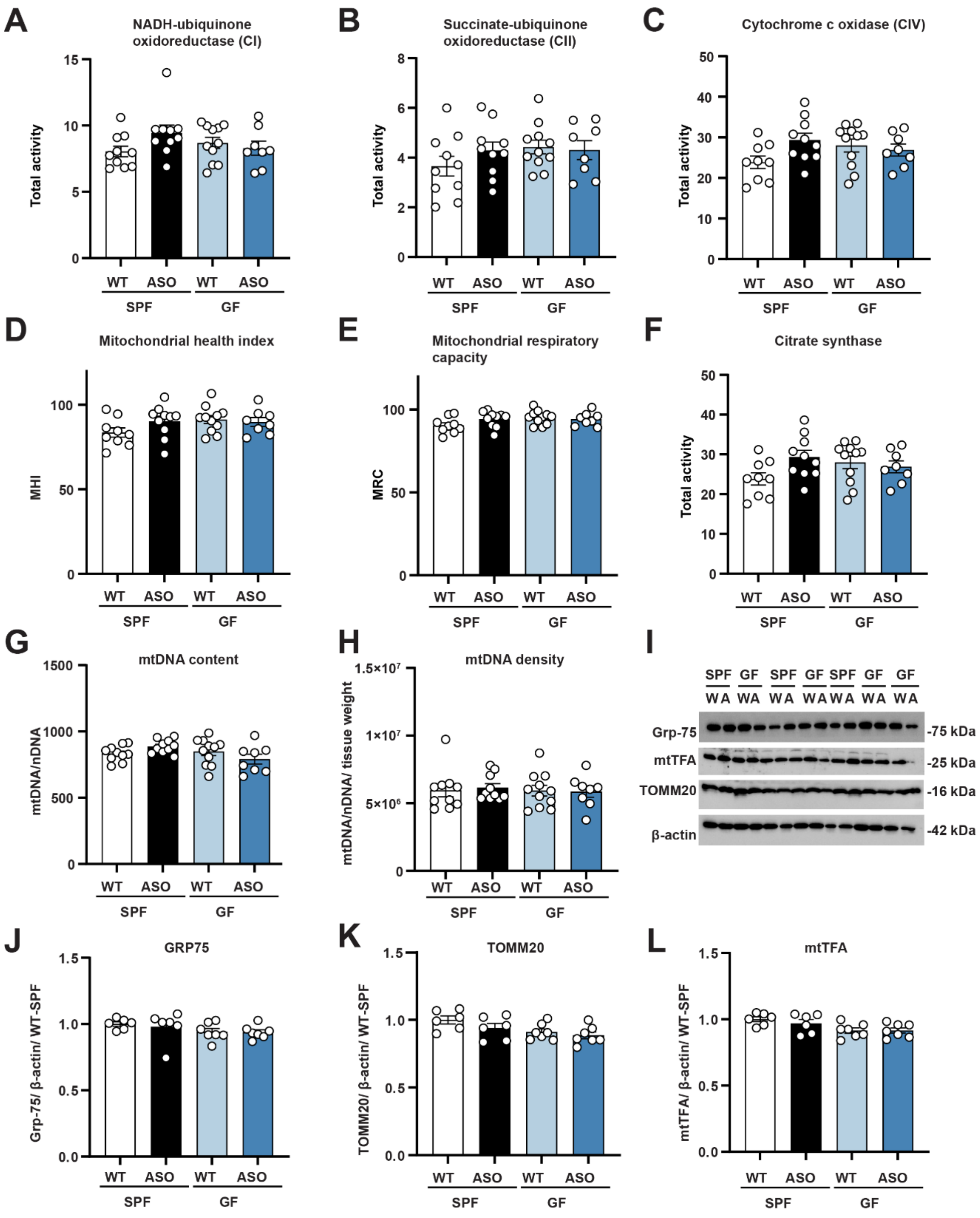
A
NADH-ubiquinone oxidoreductase (CI)
Total activity
B
Succinate-ubiquinone oxidoreductase (CII)
C
Cytochrome c oxidase (CIV)
D
Mitochondrial health index
MHI
E
Mitochondrial respiratory capacity
MRC
F
Citrate synthase
G
mtDNA content
mtDNA/nDNA
H
mtDNA density
mtDNA/nDNA/ tissue weight
I
Grp-75
mtTFA
TOMM20
β-actin
-75 kDa
-25 kDa
-16 kDa
-42 kDa
J
GRP75
Grp-75/ β-actin/ WT-SPF
K
TOMM20
TOMM20/ β-actin/ WT-SPF
L
mtTFA
mtTFA/ β-actin/ WT-SPF
WT
ASO
SPF
GF

**Fig. S3. Mitochondrial abundance is not altered by the gut microbiome or by genotype.**
(**A-C**) Mitochondrial enzymatic activity in total striatum tissue extracts for (**A**) NADH-ubiquinone oxidoreductase (CI), (**B**) succinate-ubiquinone oxidoreductase (CII), (**C**) cytochrome c oxidase (CIV). (**D**) Mitochondrial health index (MHI), calculated using the equation MHI = (CI + CII + CIV) / (CS + mtDNA density + 1) 100. (**E**) Mitochondrial respiratory capacity (MRC), calculated using the equation MRC = (Average √CI, √CII, √CIV) / (Average 3√CS, 3√mtDNA density) 100. (**F-H**) Mitochondrial content in total striatum extracts. (**F**) Citrate synthase activity. (**G**) mtDNA quantification in total striatum. (**H)** mtDNA copy number measured by qPCR, subtracting mtDNA Ct from average nDNA Ct, calculated by 2^(ΔCt) x 2. (**H**) mtDNA density, linearizing Ct as 2^Ct / (1/10^(-12)) to derive mtDNA abundance per unit of tissue. (**I-L**) Western blotting for mitochondrial markers in total striatum protein extracts. (**I**) Image of representative blot. Relative levels of (**J**) GRP75, (**K**) TOMM20, and (**L**) mtTFA in different groups. Protein levels were normalized to β-actin. SPF, specific pathogen-free; GF, germ-free; WT/W, wild-type; ASO/A, Thy1-α-synuclein overexpressing; CI, Complex I; CII, Complex II; CIV, Complex IV; CS, citrate synthase; Ct, cycle threshold; GRP75, glucose-regulated protein 75; TOMM20, translocase of outer mitochondrial membrane; mtTFA, mitochondrial transcription factor A.

**Statistical details:** (**A-H**) Data were analyzed using a linear model (variable ~ Genotype + Microbiome + Genotype*Microbiome*)* and pairwise comparisons with Benjamini-Hochberg (FDR) correction. Data are expressed as mean ± SEM. Sample size: n=6-7, data combined from 2 different cohorts. (**J-L**) Data were analyzed using a linear model (variable ~ Genotype + Microbiome + Genotype*Microbiome) and pairwise comparisons with Benjamini-Hochberg (FDR) correction. Data are expressed as mean ± SEM. Sample size: n=9-11, data combined from 2 different cohorts.

### FIGURE S4

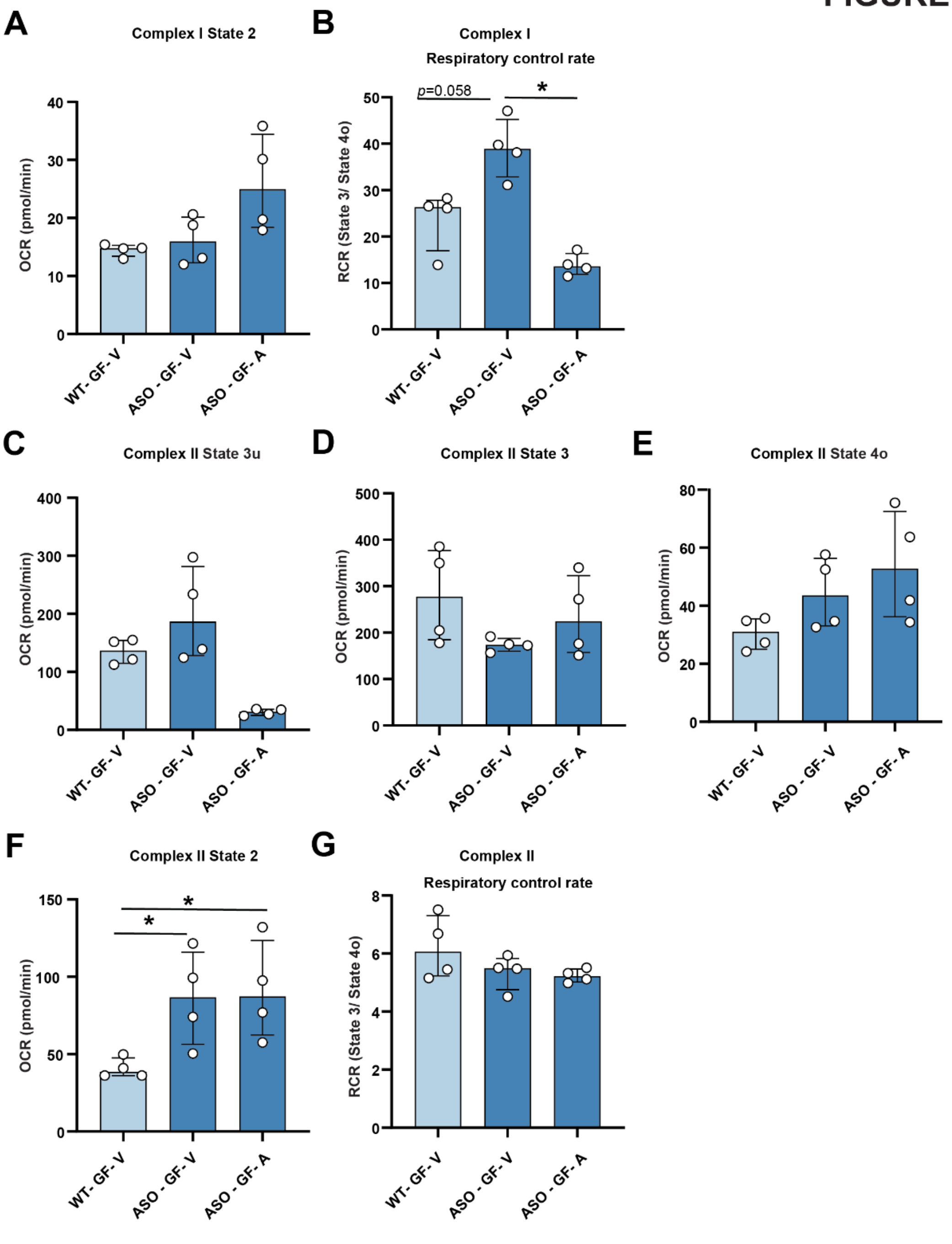
A
Complex I State 2
OCR (pmol/min)
0
10
20
30
40
WT- GF- V
ASO - GF- V
ASO - GF- A
B
Complex I
Respiratory control rate
RCR (State 3/ State 4o)
0
10
20
30
40
50
p=0.058
*
C
Complex II State 3u
0
100
200
300
400
D
Complex II State 3
0
100
200
300
400
500
E
Complex II State 4o
0
20
40
60
80
F
Complex II State 2
0
50
100
150
*
*
G
Complex II
Respiratory control rate
0
2
4
6
8

**Fig. S4. TXNRD inhibition does not affect Complex II activity in ASO-GF mice.**

(**A-G**) Oxygen consumption rate (OCR) was measured in freshly isolated mitochondria using Seahorse, evaluating Complex I (CI) and Complex II (CII) respiration. (**A**) CI State 2. (**B**) CI respiratory control rate (RCR). (**C**) CII State 3u (uncoupled, maximal respiration). (**D**) CII State 3 (ATP-linked respiration). (**E**) CII State 4o (proton leak). (**F**) CII State 2. (**G**) CII RCR. SPF, specific pathogen-free; GF, germ-free; WT, wild-type; ASO, Thy1-α-synuclein overexpressing; V, vehicle; A, Auranofin.

**Statistical details:** (**A-G**) Data were analyzed by Kruskal-Wallis test followed by Dunn's post-hoc test with Benjamini-Hochberg (FDR) correction. Significance: * $p < 0.05$. Data are expressed as median ± interquartile range. Data combined from 2 different cohorts. Sample size: n=4 per group.

**Table S2. Key Resource Table**
Data, code, protocols, and key lab materials used in this study.

| RESOURCE TYPE | RESOURCE NAME | SOURCE | IDENTIFIER | NEW/ REUSE | ADDITIONAL INFORMATION |
|---|---|---|---|---|---|
| Dataset | Tabular behavior data | GitHub | https://github.com/jdthoang/LHMorais2024 | new | data from pole descent and beam traversal assays shown in Figures 1A-B and 4A-D |
| Dataset | Tabular mitochondrial function data | GitHub | https://github.com/jdthoang/LHMorais2024 | new | data from mitochondrial function profiling shown in Fig. 2, Fig. 4E-G, S2, S3, and S4 |
| Dataset | Transcriptomics data | GitHub | https://github.com/jdthoang/LHMorais2024 | new | data from transcriptomics shown in Fig. 1C-G and S1 |
| Dataset | Proteomics data | GitHub | https://github.com/jdthoang/LHMorais2024 | new | data from proteomics shown in Fig. 3A-F |
| Software/code | Analysis and visualization code | GitHub | https://github.com/jdthoang/LHMorais2024 | new | code used to analyze all data |
| Software/code | Rsubread v2.16.1 | https://bioconductor.org/packages/release/bioc/html/Rsubread.html | RRID:SCR_016945 | reuse | |
| Software/code | limma v3.60.4 | https://bioconductor.org/packages/release/bioc/html/limma.html | RRID:SCR_010943 | reuse | |
| Software/code | DESeq2 v1.42.1 | https://bioconductor.org/packages/release/bioc/html/DESeq2.html | RRID:SCR_015687 | reuse | |
| Software/code | RITAN v1.26.0 | https://bioconductor.org/packages/release/bioc/html/RITAN.html | N/A | reuse | |
| Software/code | Xcalibur | https://www.thermofisher.com/order/catalog/product/OPTON-30965 | RRID:SCR_014593 | reuse | |
| Software/code | Proteome Discoverer v2.5 | https://www.thermofisher.com/order/catalog/product/IQLAAEGABSFAKJMAUH | RRID:SCR_014477 | reuse | |
| Software/code | UniProtKB | http://www.uniprot.org/help/uniprotkb | RRID:SCR_004426 | reuse | |
| Software/code | Sequest with Percolator | http://noble.gs.washington.edu/proj/percolator/ | N/A | reuse | |
| Software/code | MitoCarta v3.0 | http://www.broadinstitute.org/pubs/MitoCarta/ | RRID:SCR_018165 | reuse | |
| Software/code | Python v3.11.5 | http://www.python.org | RRID:SCR_008394 | reuse | |
| Software/code | NumPy v1.24.3 | http://www.numpy.org | RRID:SCR_008633 | reuse | |
| Software/code | Pandas v2.1.1 | https://pandas.pydata.org | RRID:SCR_018214 | reuse | |
| Software/code | Numba v0.58.0 | https://numba.readthedocs.io/en/stable/index.html | RRID_SCR_025056 | reuse | |
| Software/code | Bokeh v3.2.0 | https://bokeh.org | N/A | reuse | |

| | | | | | |
|---|---|---|---|---|---|
| Software/code | bebi103 v0.1.17 | https://bebi103.github.io | N/A | reuse | |
| Software/code | JupyterLab v4.0.6 | http://jupyterlab.github.io/jupyterlab/ | RRID:SCR_023339 | reuse | |
| Software/code | R v4.3.2 | https://www.r-project.org/ | RRID:SCR_001905 | reuse | |
| Protocol | Pole descent | protocols.io | dx.doi.org/10.17504/protocols.io.n92ld8j87v5b/v1 | reuse | |
| Protocol | Beam traversal | protocols.io | dx.doi.org/10.17504/protocols.io.e6nvw1212lmk/v1 | new | |
| Protocol | Tissue dissection | protocols.io | dx.doi.org/10.17504/protocols.io.x54v92z2ql3e/v1 | new | |
| Protocol | Oxidative stress quantification | protocols.io | dx.doi.org/10.17504/protocols.io.4r3l2qbqjl1y/v1 | new | |
| Protocol | Mitochondrial isolation | protocols.io | dx.doi.org/10.17504/protocols.io.5jyl82qz6l2w/v1 | new | |
| Protocol | Mitochondrial respirometry | protocols.io | dx.doi.org/10.17504/protocols.io.bp2l62z4zgqe/v1 | new | |
| Protocol | Western blotting | protocols.io | dx.doi.org/10.17504/protocols.io.x54v92z2ql3e/v1 | new | |
| Protocol | Immuno-histochemistry | protocols.io | dx.doi.org/10.17504/protocols.io.kqdg32y61v25/v1 | new | |
| Protocol | Mitochondrial proteomics | protocols.io | dx.doi.org/10.17504/protocols.io.36wgqny15gk5/v1 | new | |
| Antibody | Rabbit anti-TFAM | Abclonal | Cat# A1926, RRID:AB_2763953 | reuse | 1:500 dilution |
| Antibody | Donkey anti-Rabbit IgG H&L Alexa Fluor 647 | Abcam | Cat# ab150075, RRID:AB_2752244 | reuse | 1:500 dilution |
| Chemical, peptide, or recombinant protein | N-acetyl-L-cysteine | Sigma-Aldrich | Cat. #A7250 | reuse | |
| Chemical, peptide, or recombinant protein | Auranofin | Cayman | Cat. #15316 | reuse | |
| Critical commercial assay | OxiSelect In Vitro ROS/RNS Assay Kit | CellBio Labs | Cat. #STA-347-5 | reuse | |
| Experimental model: Organism/strain | "Line 61" Thy1-αSyn mice | Masliah Lab | N/A | reuse | Chesselet et al. Neurother 9, 297–314 (2012); Rockenstein et al J Neurosci Res 68:568–578 (2002) |
| Experimental model: Organism/strain | BDF1 mice | Charles River | RRID:IMSR_CRL:099 | reuse | |

| | | | | | |
|---|---|---|---|---|---|
| Oligonucleotide | COX1-Fwd | Rosenberg et al. Nat Commun 14, 4726 (2023) | N/A | reuse | Sequence: ACCACCATCATTTCTCCTTCTC |
| Oligonucleotide | COX1-Rev | Rosenberg et al. Nat Commun 14, 4726 (2023) | N/A | reuse | Sequence: CTCCTGCATGGGCTAGATTT |
| Oligonucleotide | COX1-Probe | Rosenberg et al. Nat Commun 14, 4726 (2023) | N/A | reuse | Sequence: HEX/AAGCAGGAG/ZEN/CAGGAACAGGATGAA/3IABkFQ |
| Oligonucleotide | B2M-Fwd | Rosenberg et al. Nat Commun 14, 4726 (2023) | N/A | reuse | Sequence: GAGAATGGGAAGCCGAACATA |
| Oligonucleotide | B2M-Rev | Rosenberg et al. Nat Commun 14, 4726 (2023) | N/A | reuse | Sequence: CCGTTCTTCAGCATTTGGATTT |
| Oligonucleotide | B2M-Probe | Rosenberg et al. Nat Commun 14, 4726 (2023) | N/A | reuse | Sequence: FAM/CGTAACACA/ZEN/GTTCCACCCGCCTC/3IABkFQ |
